# Multivariate analysis of glycogenes reveals coordinated regulation of immunoglobulin glycosylation in an immortalized human B cell system

**DOI:** 10.1101/2025.09.30.679651

**Authors:** Christine D. Wiggins, Marina Sangés Ametllé, Claire Wu, Timothy A. Watkins, Kristin L. Boswell, Richard A. Koup, Douglas A. Lauffenburger

**Affiliations:** Department of Biological Engineering, Massachusetts Institute of Technology, Cambridge, MA 02142, USA; Technical University of Denmark, Copenhagen, Denmark; Vaccine Research Center, National Institute of Allergy and Infectious Diseases, National Institutes of Health, Bethesda, MD, United States

## Abstract

While neutralizing ability has traditionally been considered the most important antibody function, appreciation has grown for Fc-mediated ‘extra-neutralizing’ functions, which are shaped by IgG glycosylation. However, there remain fundamental questions as to how B lymphocytes induce and regulate antibody glycosylation and thus functional capability. Understanding how transcriptional and cell state regulation shape glycosylation could reveal levers to tune protective humoral profiles in a disease- and antigen-specific manner. Prior studies have explored a limited panel of glycogenes and measured bulk glycosylation changes. Here, employing an *in vitro* antigen-specific B cell culture system, we systematically characterize transcriptional and humoral responses to cytokine perturbations. After exposure to a broad panel of cytokines (IL-4, IL-6, IL-10, IL-17, TNFa, IFNg, APRIL, and BAFF) across multiple concentrations and timepoints, transcriptomic profiling and lectin-based IgG glycome assays are employed to associate cytokine stimuli with both glycogene expression and IgG glycosylation. Supervised and unsupervised machine learning models identify cytokine-specific glycogene “signatures” as well as distinct immunoglobulin glycosylation profiles. We find that cytokines induce rapid transcriptional responses, with glycogene signatures outperforming single-gene changes in distinguishing stimulation conditions. We further demonstrate the ability to induce both pro- and anti-inflammatory IgG glycosylation profiles, particularly in terms of IgG galactosylation. This work demonstrates the utility of this system to parse the cytokine-driven regulation of B lymphocyte glycogenes, establishing a framework for dissecting how environmental cues shape antibody glycosylation, with relevance for autoimmune disease, infection, and vaccine responses.

## INTRODUCTION

Antibodies are glycoproteins produced by B cells in response to activation by a pathogen. Glycan structures on the fragment-crystallizable (Fc) region influence an antibody’s interaction with Fc gamma receptors (FcgRs) present on innate immune cells, thereby modulating downstream functional immune responses such as antibody-dependent cellular cytotoxicity (ADCC), complement deposition (ADCD), cellular phagocytosis (ADCP), and natural killer cell activity (ADNK)^1–3^. The reliance of these downstream functions on Fc binding and glycosylation status is well-studied, with glycoengineering studies *in vitro* demonstrating 17-fold higher binding to FcgRIIIa and FcgRIIIb in hypofucosylated IgG1, leading to enhanced NK cell-mediated ADCC as well as C1q binding^4^. Further studies have linked afucosylation and hypergalactosylation with enhanced ADCC^5^. Meanwhile, antibody sialylation has been linked to therapeutic anti-inflammatory effects, potentially through decreased affinity for type 1 Fc receptors and reduced IgG-driven ADCC, though precise mechanisms are disputed^6^. These differences are clinically relevant-alterations in antibody glycosylation and resulting changes in functional responses have been noted in the context of infection, vaccination, autoimmunity, pregnancy, and aging, though direct mechanistic causes have been challenging to find^7–15^.

Glycosylation is thought to be regulated at multiple levels, with altered IgG glycosylation linked to changes in transcription factor levels, glycogene expression, hormone and cytokine concentrations, and other biological factors^16–21^. In many contexts where altered glycosylation has been observed, cytokines are present in complex mixtures, and their direct links to antibody glycosylation are further obfuscated by patient heterogeneity and temporal considerations. Altered levels of cytokines such as IL-4, IL-10, IL-6, IL-17, IFNg, or TNFa have been observed in autoimmune conditions such as systemic lupus erythematosus (SLE) and rheumatoid arthritis (RA), as well as in the context of severe infection with viruses such as SARS-CoV-2 or influenza, all conditions where altered glycosylation has also been observed^22–26^. As cytokines are key regulators of B cell proliferation, survival, metabolism, and class switching, their role in influencing IgG glycosylation is of interest and remains poorly defined. Various approaches have been utilized *in vitro* to understand alterations in antibody glycosylation due to cytokine milieu. Studies have linked cytokine stimulation with altered bulk IgG glycosylation, but most have tested only single concentrations of a handful of cytokines, considered only one stimulation timepoint, or not considered combinatorial cytokine stimulations^19,20^.

Further, comprehensive transcriptomic profiling has not been explored in this context. While RT-PCR studies have captured specific glycosyltransferase changes due to cytokine supplementation, suggesting cytokine regulation of gene-level control over glycosylation, up to 5% of the human transcriptome, termed “glycogenes”, may be implicated in glycan structure and function, necessitating deeper transcriptomic exploration^27^. As regulation of glycosylation is polygenic, univariate analysis of gene regulation moreover risks missing the complex interplay of multiple glycogenes required to form any particular type of glycosylation. Additionally, antigen-specific glycosylation patterns have been observed to diverge from bulk glycan status, while existing *in vitro* studies of cytokine influence have primarily explored bulk glycan status^28,29^. Knowledge of combinatorial or concentration-dependent cytokine influences on gene-level regulation of glycosylation of antibodies in an antigen-specific manner could inform the use of cytokine-modifying treatments in infectious disease or affect vaccine design strategies in a bid to leverage the downstream functional implications of altered antibody glycosylation.

Though efforts have been made to explore the role of environmental parameters such as cytokine stimulation in antibody glycosylation, existing model systems have significant limitations. Many models are not human-derived or fully B-cell derived, relying on experimentally tractable cell lines such as Chinese hamster ovary (CHO) cell lines^30–32^. Species and cell-type specific regulation of glycosylation reduces the human *in vivo* relevance of any findings in systems of these types. Other model systems have attempted to use primary human B cells, which avoid the issues of species and cell-type specific regulation mismatches, but the short lifespan of primary cells combined with the low frequency of antigen-specific B cells in serum even directly after vaccination limits systematic study^33,34^. To explore glycosylation regulation in an antigen-specific system, only a small number of stimulation conditions and timepoints would be feasible, as cell number constraints would be very limiting. Even without antigen-specific stimulation, IgG produced by antigen-specific B cell clones varies in glycosylation from IgG produced by donor-matched control B cell clones, suggesting that IgG glycosylation differences observed in antigen-specific systems may be different from those observed in bulk systems more commonly utilized^35^.

Here, we employ an antigen-specific, immortalized human B cell model to systematically study cytokine regulation of both IgG glycosylation and glycogene expression. This system provides us with a manipulable model that allows for controlled cytokine exposures of multiple timescales and concentrations, with both transcriptomic and glycomic readouts. We profile cytokine-driven changes at the transcriptome and glycome level, using multivariate machine learning methods to identify cytokine-specific glycogene and glycosylation patterns.

## RESULTS

### Bimms sustain proliferation over time, with appropriate stimulation responses and continuous immunoglobulin secretion

To investigate the role of cytokine stimulation in immunoglobulin glycosylation, we employed a co-culture system, with pools of H7 HA-specific IgG+ memory B cells immortalized via *BCL6*, *BCLXL*, and GFP overexpression through retroviral transduction^36,37^. When grown in the presence of IL-21 on irradiated feeder cells expressing CD40L (Fig 1A), the cell lines have been shown to proliferate for at least 6 months, possess a low mutation rate, secrete immunoglobulin, and have signaling-capable B cell receptors (BCRs)^36,38^. 3 IgG-secreting immortalized B cell clonal lines (Bimms) from each of 2 donors, F2 and F5, were selected and pooled by donor (Fig 1B). With this approach, experimental dependence on any one clone is reduced while still allowing for the incorporation of potential donor-to-donor variability. However, individuals produce many antigen-specific clonal B cells in response to vaccination or infection, and so these Bimm pools should not be thought of as representative of an individual’s full B cell response. All clones included in Bimm pools proliferated over the course of 21 days, with most lines reaching peak cell density by day 18 (Fig 1C). For irradiated YK6 feeder cells, no cell proliferation was seen over the course of 10 days of culture, with cell death occurring after day 7 (Supp Fig 1A). Therefore, for all Bimm cultures, YK6 feeder cells were supplemented or replaced by day 7. Flow cytometry shows the ability of media rinsing followed by CD19 positive selection via magnetic activated cell sorting (MACS) to provide 98.6% (n = 5) Bimm purity for downstream analysis, with YK6 content (Supp Fig 1B) decreasing from trypsin-lifted co-cultures (Supp Fig 1C) by media rinsing alone (Supp Fig 1D) or followed by MACS (Supp Fig 1E).

**Figure 1.**
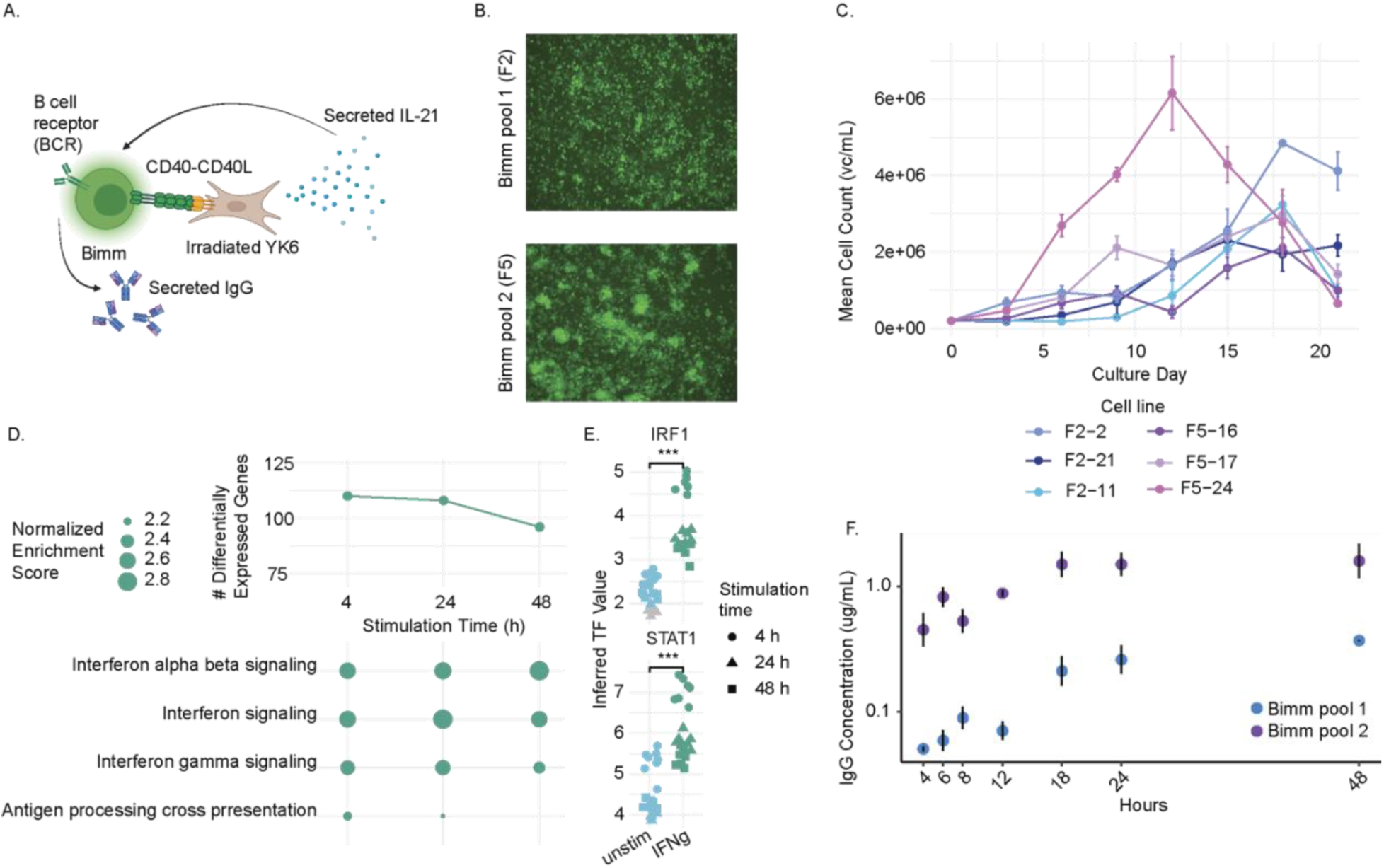
Immortalized B cells proliferate, respond to stimuli, and secrete immunoglobulin. A) Schematic of interactions between Bimms and irradiated feeder cells. B) GFP images of unstimulated Bimms. C) Growth curves for each of 6 Bimm cell lines, n = 3 replicates. Mean +/− 1 standard deviation shown. D) Number of differentially expressed genes (p.adj < 0.05, abs(log2 fold change) > 1) and Reactome gene set enrichment for IFNg stimulated samples across 4, 24, and 48 hours of stimulation. Size of dot for gene set enrichment corresponds to normalized enrichment score value. E) Inferred transcription factor values for IRF1 and STAT1 for unstimulated or IFNg-stimulated samples at 4, 24, or 48 hours. Inferred values are colored by significance (grey, inferred at p > 0.05) and condition (unstimulated = blue, IFNg stimulated = green). Statistical significance between stimulations was performed via Mann-Whitney U test (*** p < 0.001). F) Media concentration of total IgG over time per Bimm pool measured via total human IgG ELISA, shown as mean +/− 1 standard deviation. Samples measured in triplicate, n = 3 biological replicates.

Since Bimm purity was sufficient for RNAseq analysis, we sought to confirm that cytokine stimulation would induce transcriptional changes. We stimulated one Bimm clone with IFNg alone, a-IgG for BCR stimulation, or in combination. As no genes were differentially expressed at any timepoint between BCR-stimulated or unstimulated cells (Supp Fig 1F-H), all IFNg-stimulated samples, regardless of BCR stimulation, were grouped. IFNg stimulation induced differential gene expression as early as 4 hours post-stimulation (Fig 1D), with the number of differentially expressed genes (DEGs) decreasing slightly by 48 hours of stimulation. Many of these differentially expressed genes code for genes known to be induced by IFNg stimulation, such as *STAT1, GBP5, GBP2, CXCL9,* and *CD38* (Supp Fig 1I)^39,40^. Reactome pathway enrichment revealed that, of DEGs present at 2 or more timepoints, many of them belong to classic interferon signaling pathways (Fig 1D).

As several of the DEGs across timepoints were coding for transcription factors such as STAT1 (Supp Fig1I-K), we next inspected the inferred transcription factor (TF) activity predicted from gene expression data using CollecTRI regulons with the decoupleR algorithm^41^. DecoupleR allows for the inference of TF activity from transcriptomics data, utilizing prior knowledge of TF control of gene transcription contained within the CollecTRI regulons. We specifically looked at predicted TF activity for TFs known to be regulated in B cells by IFNg such as IRF1 and STAT1, comparing activity between unstimulated and IFNg stimulated conditions. Considering only TF activity inferred to be significant, IRF1 and STAT1 showed significantly higher activity in IFNg stimulated cultures compared to unstimulated, with the highest activity inferred at 4 hours of stimulation for both. This result suggests that IRF1 and STAT1 are active TFs regardless of stimulation, with the magnitude of activity significantly amplified after IFNg stimulation, aligning with previous findings^42,43^. Additionally, temporal dynamics are observable, with 4 hours of stimulation inducing more IRF1 and STAT1 activity than 24 or 48 hours. These data suggest that Bimms are capable of transcriptional responses to cytokines, in ways that align with prior knowledge. Further, Bimms exhibit time-dependent responses to IFNg, underscoring their ability to perform complex temporal signaling.

Finally, we sought confirmation that these Bimm pools could produce and secrete immunoglobulin at a detectable level. We measured IgG in media supernatant over the course of 48 hours using a total human IgG enzyme-linked immunosorbent assay (ELISA). Both Bimm pools secreted detectable IgG into the supernatant above the ELISA limit of quantification, with close agreement between replicates (Fig 1F). Bimm pool 2 overall secretes more IgG than Bimm pool 1, but both pools showed a similar trend of increasing supernatant IgG concentration over time. Thus, both Bimm pools were judged suitable for characterization of immunoglobulin after stimulation.

### Short-term cytokine stimulation induces transcriptional changes

Cytokine stimulation is known to influence both antibody glycosylation and expression of several glycogenes. However, studies to this point have focused on a single concentration of cytokine and have been restricted in ability to perform untargeted RNAseq by cell number constraints^19,20^. We instead aimed to take a systems approach to understanding the relationship between cytokine stimulation and glycogene expression. We selected 8 cytokines, IL-4, IL-6, IL-10, IL-17, TNFa, IFNg, APRIL, and BAFF, previously shown to be associated with B cell-related autoimmune conditions such as systemic lupus erythromatosus (SLE), produced by T follicular helper cells (Tfh) in the germinal center, or associated with altered IgG glycosylation *in vitro* in small scale experiments^19,24,44^. We selected two concentrations of each cytokine, a high and a low concentration, based on previous findings on the role of cytokine concentration on IgG production in an antibody-secreting cell (ASC) culture system (Table 1)^19^. Further, as Bimms have signaling-capable BCRs, we also included as a condition BCR cross-linking via a-IgG antibody. Overall, this amounted to 36 stimulation conditions applied to each of our two Bimm pools (Fig 2A).

**Figure 2.**
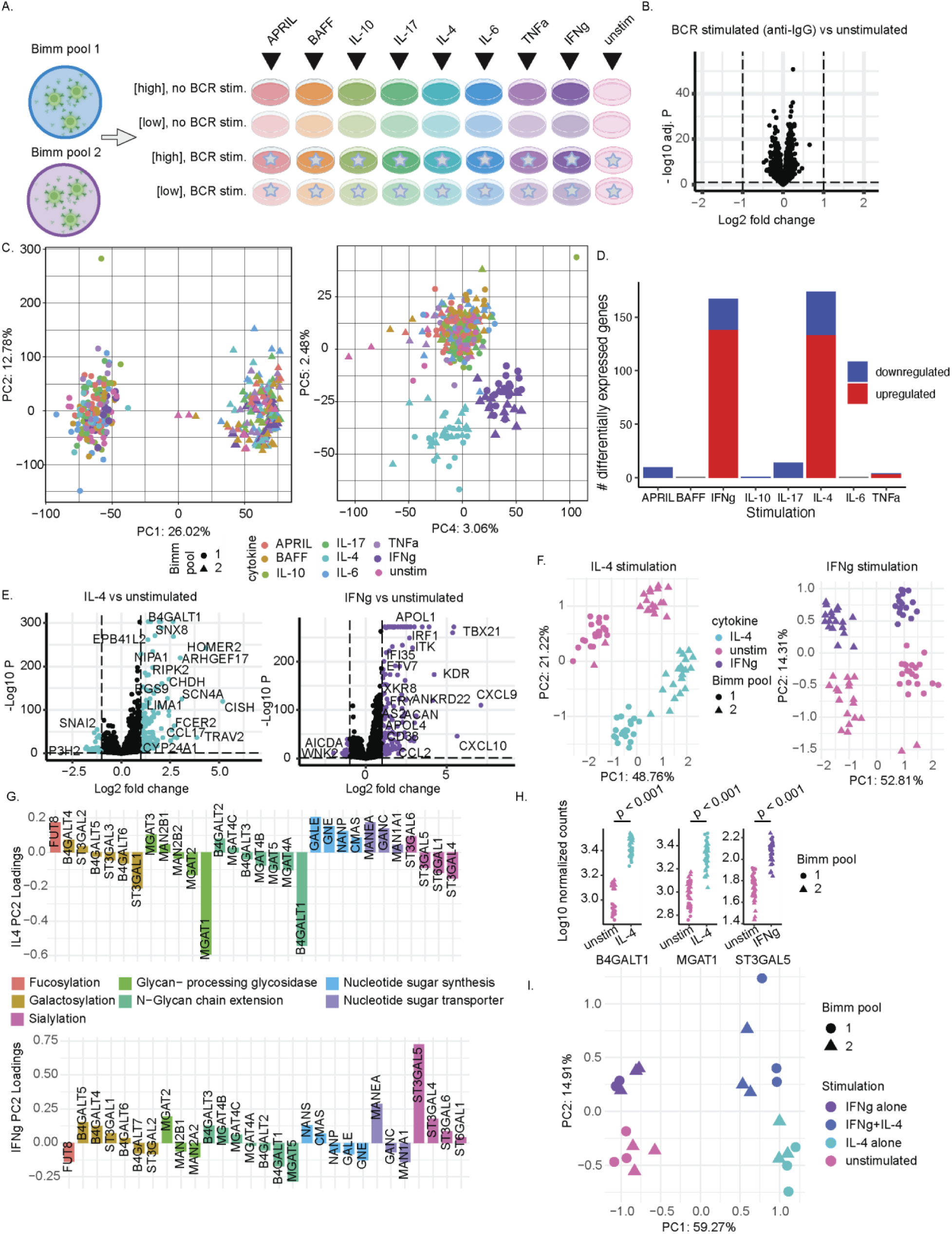
IL-4 and IFNg drive glycogene changes solo or in combination. A) Schematic of stimulation experiment, with 8 cytokines supplemented at either a high or low concentration, both with and without BCR stimulation, for two Bimm pools. B) Differentially expressed genes for a BCR stimulated condition vs unstimulated, adjusted p < 0.05, |log2 fold-change| > 1. C) Principal components scores plot for all genes on principal components 1 and 2 (left) and 4 and 5 (right). D) Number of differentially expressed genes per stimulation condition. E) Differentially expressed genes for IL-4 (left) and IFNg (right) conditions compared to unstimulated controls, adjusted p < 0.05, |log2 fold-change| > 1. F) Principal components scores plot for known glycogenes in IL-4 stimulation (left) and IFNg stimulation (right). G) Principal components loadings for PC2 for IL-4 (top) and IFNg (bottom) PCAs, with loadings colored by the glycogene family they belong to. Only loadings with |value| > 0.02 are shown. H) Log10 normalized counts for glycogenes statistically significantly different from unstimulated controls in either IL-4 or IFNg stimulated conditions. Statistical significance assessed via DESeq2. I) Principal components scores plot for glycogenes from combinatorial stimulation with IL-4 and IFNg.

Initially, we selected a 4 hour stimulation timepoint to investigate immediate signaling changes upon exposure to relevant cytokines, as strong transcriptional differences were apparent at this time with IFNg stimulation. Bimm pools were grown for 14 days prior to stimulation with each of 36 unique stimulation conditions. As no genes were differentially expressed between BCR-stimulated or unstimulated cells (Fig 2B) across conditions, all samples were grouped by cytokine stimulation, regardless of BCR stimulation. We hypothesize that overexpression of BCL6 for the purposes of cell line immortalization may be masking any expected transcriptional changes from BCR stimulation, as BCL6 is a known repressor of IRF4, through which much of the signaling cascade initiated by BCR stimulation flows^45^. Further, at the 4 hour timepoint, no genes were differentially expressed between any high and low cytokine concentration, and so both cytokine stimulation levels were analyzed together.

Principal component analysis performed with all genes revealed high variability between Bimm pools (Fig 2C), with Reactome pathway enrichment demonstrating that much of the variability between pools is tied to differences in cell growth, cell-cell junctions, and transcription (Supp Fig2A). These findings align with the differences seen morphologically between Bimm pools (Fig 1B). While most variability in RNAseq captured by PCA is tied to pool differences, higher-order PCs such as PCs 4 and 5 capture variability driven by stimulation condition, particularly demonstrating strong changes tied to IL-4 and IFNg stimulation (Fig 2C). These differences appear to be consistent between Bimm pools, and result in samples distinct from other stimulation conditions.

Looking at differentially expressed genes between stimulated and unstimulated conditions, we see that IFNg and IL-4 do indeed induce the largest number of DEGs, primarily genes whose expression is upregulated during stimulation (Fig 2D). In IL-4 stimulation, genes such *HOMER2, ARGHEF17, CISH*, *CCL17*, and *FCER2* are expectedly upregulated (Fig 2E)^46,47^. Interestingly, one of the most significantly upregulated genes, *B4GALT1*, encodes a galatosyltransferase that plays a key role in N-glycosylation^15^. In IFNg stimulation, expected genes such as *CD38*, *CXCL9*, *CXCL10*, and *IRF1* are upregulated^40^.

We then selected a panel of 42 genes shown to be involved in various IgG N-glycosylation processes and performed PCA on these genes (Table 2, Supp Fig 2B)^48^. IL-4 stimulated conditions from both Bimm pools strongly separated from all other conditions, and PCA scores plots on individual stimulations reveal that these glycogenes do indeed vary strongly between both stimulated and unstimulated conditions as well as Bimm pools for both IL-4 and IFNg stimulation (Fig 2F). Interestingly, while unstimulated Bimm pools start in different locations in this reduced dimensionality glycospace, stimulation by either IL-4 or IFNg moves each pool in a comparable glycospace direction, suggesting a shared common response to stimulation regardless of initial starting point. To examine the glycogene changes responsible for this shared response, we can examine the PCA loadings. Loadings for a PCA should be thought of as a comprehensive signature, where the combination of contributions from each glycogene is what differentiates samples from different conditions. PC1 loadings of genes with high (|loading value| > 0.02) loadings for both IL-4 and IFNg scores plots show very similar trends for signatures tied to Bimm pool separation regardless of stimulation, with *ST3GAL6*, *B4GALT1*, *B4GALT2*, *B4GALT5*, and *FUT8* expression higher in Bimm pool 1, and glycogenes such as *MGAT5, MGAT4A, ST6GAL1, B4GALT6,* and *ST3GAL1* higher in Bimm pool 2 (Supp Fig 2C, 2D). As nearly 50% of glycosylation differences have been linked to genetic heritability, these distinct starting locations for each Bimm pool make sense^49^. Examining the PC2 loadings with high (|loading value| > 0.02) loadings describing separation by stimulation, we see that for IL-4 stimulation, the largest contributors to IL-4 scores are *MGAT1* and *B4GALT1*, but other genes associated with sialylation such as *ST3GAL5, ST6GAL1,* and *ST3GAL4* are also higher in IL-4-stimulated samples (Fig 2G). For IFNg loadings, *ST3GAL5* is the highest contributor to separation and is involved in sialylation, but other sialylation-related genes such as *ST3GAL5* also play a part in the signature, as do sugar transporters such as *MANEA*. Unsurprisingly, the highest contributors to the PCA glycogene signature are also univariately significantly overexpressed in stimulated conditions across both Bimm pools (Fig 2H).

### IL-4 and IFNg stimulation exert additive effects on glycogenes

Because IL-4 and IFNg exposure both have large effects on glycogene expression, we next sought to determine if those effects were opposing, complimentary, or additive. We stimulated Bimm pools for 4 hours with either IFNg alone, IL-4 alone, blank media, or a combination of IFNg and IL-4. We found again that cells treated with IFNg expressed classic IFNg-controlled genes such as *GBP5*, *IRF1, STAT1, CXCL9* among others when compared to cells with no IFNg treatment (Supp Fig 2E, 2F). In IL-4-containing conditions, *FCER2, HOMER2,* and *ARHGEF17* were again upregulated (Supp Fig 2E, 2G). The dual-stimulated condition then expectedly overexpressed a combination of these most highly differentially expressed genes. Building a PCA of all DEGs, samples separated neatly by stimulation condition, with the majority of variance coming from stimulation and not Bimm pool (Supp Fig 2H). A PCA scores plot focusing specifically on glycogenes demonstrates the combinatorial effect of IFNg and IL-4, with PC1 describing the IL-4 stimulation effect, PC2 describing the IFNg stimulation effect, and the combinatorial samples found in a quadrant suggesting an additive effect (Fig 2I). The glycogenes contributing most to IL-4 (PC1) and IFNg (PC2) separation were the same as in the larger experiment (Supp Fig 2I, 2J), further demonstrating the consistency of cytokine-induced glycogene changes at 4 hours of stimulation.

### Long-term stimulation reshapes transcriptional and glycosylation profiles

Many chronic disease conditions associated with altered immunoglobulin glycosylation, such as SLE, rheumatoid arthritis, and multiple sclerosis, are also characterized by long-term alterations in inflammatory status and signaling environment, so we next sought to characterize the effects of longer-term cytokine stimulation on glycosylation^24,50,51^. We repeated the previous experiment, but now with 14 days of stimulation. All conditions were included, except for IFNg, which led to widespread cell death by day 7 even at lower concentrations. Further, BCR stimulation was applied only for the first 3 days of the experiment due to increased cell death as well. For this timepoint, transcriptomic, proteomic, and glycomic readouts were taken (Supp Fig 3A). IL-4 stimulation induced a high number of differentially expressed genes as did TNFa stimulation, and IL-17 stimulation to a lesser extent (Fig 3A). Similarly to findings at 4 hours, BCR stimulation did not induce any differentially expressed genes (Supp Fig 3B).

**Figure 3.**
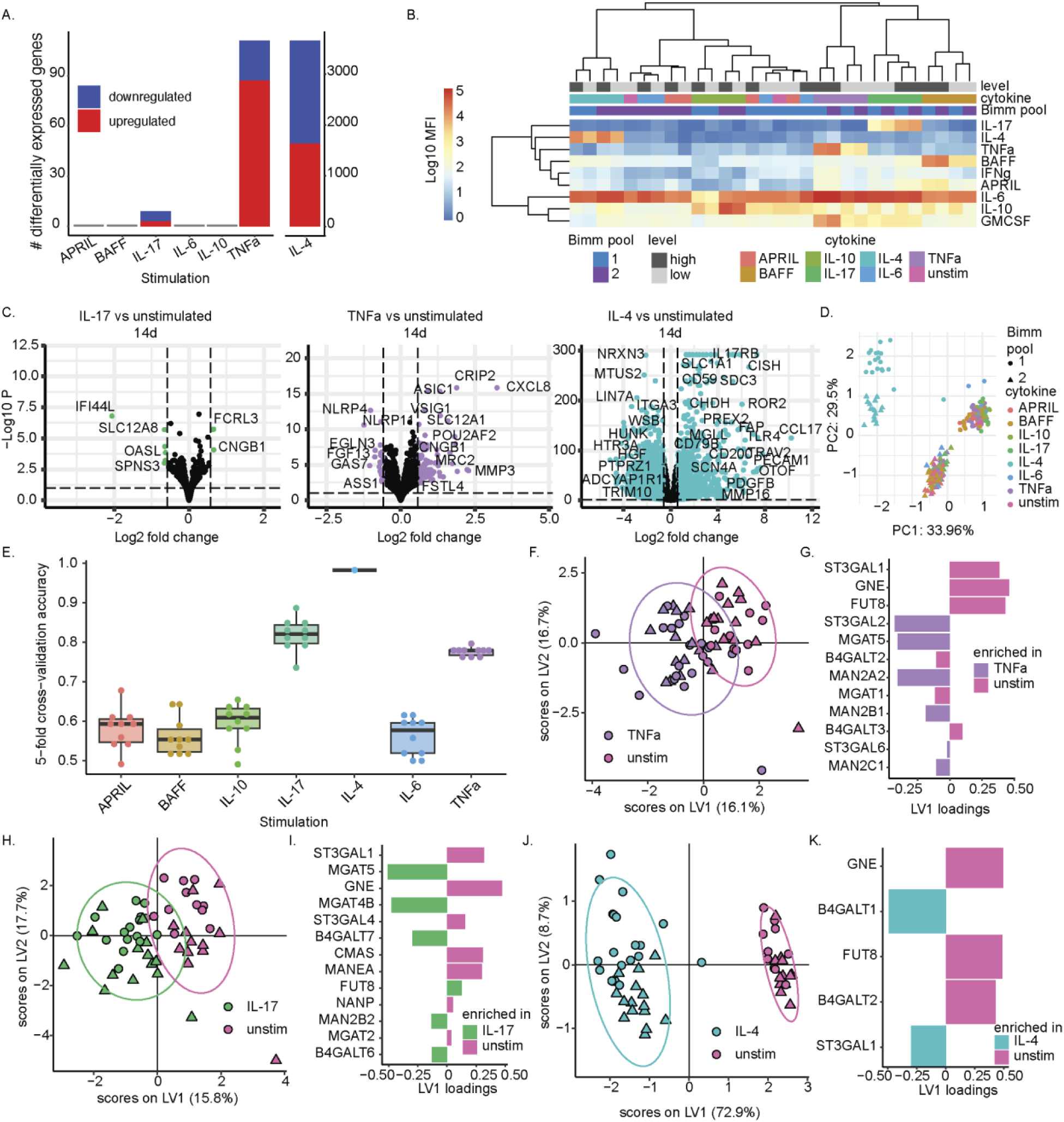
Multivariate signatures describe longer-term shifts in glycogenes. A) Number of differentially expressed genes per stimulation condition. B) Heatmap of Luminex measurements of supernatant cytokines and chemokines, averaged by Bimm pool, cytokine stimulation condition, and cytokine stimulation level. Samples are clustered via Euclidean clustering. C) Differentially expressed genes for IL17, TNFa, and IL4 conditions compared to unstimulated controls, adjusted p < 0.05, |log2 fold-change| > 0.58. D) Principal components scores plot for known glycogenes. E) 5-fold cross validation accuracy of partial least squares discriminant analysis models built on glycogenes for each pairwise comparison of stimulation to unstimulated condition. Each dot represents the one round of 5-fold cross validation, with boxplots displaying the median value, with lower and upper hinges representing the 25^th^ and 75^th^ percentiles, with the whiskers plotted between the hinges to the largest and smallest values at most 1.5x the interquartile range from their respective hinges. F) PLS-DA scores plot and G) Loadings plot for TNFa stimulation vs unstimulated controls.H) PLS-DA scores plot and I) Loadings plot for IL17 stimulation vs unstimulated controls. J) PLS-DA scores plot and K) Loadings plot for IL4 stimulation vs unstimulated controls. For all scores plots, score shape corresponds to Bimm pool (circle = Bimm pool 1, triangle = Bimm pool 2), while ellipses represent a 95% confidence interval. For loadings plots, enrichment denotes the stimulation condition with the higher mean expression of a given gene.

To understand why only some of the stimulation conditions resulted in differential gene expression, we characterized the media supernatant immediately prior to the end of the experiment for cytokine content. Using a custom 9-plex Luminex kit, we assayed the media for levels of all supplemented cytokines, plus IFNg and GMCSF. We found that supplementation did indeed lead to higher supplemented cytokine levels for IL-17, IL-4, TNFa, and BAFF, where low cytokine supplementation levels also led to lower detected cytokine in media supernatant (Fig 3B). For IL-10, higher levels of IL-10 were detected in supplemented conditions, but Bimm pool 2 experienced higher levels than Bimm pool 1 in the media by the end of the experiment regardless of supplementation level, whether due to lower consumption or possible production. IL-6, which may be produced by B cells and contributes to germinal center formation, appears to have been produced by all cells, resulting in a stimulation condition undifferentiated from both control and other stimulations^52^. Finally, APRIL stimulation does not result in enhanced APRIL levels by the end of experiment, likely due to consumption. As BAFF stimulation did result in a differentiated and stimulation-level-dependent condition, but no differentially expressed genes, we hypothesize that the process of Bimm immortalization, with overexpression of *BCL6*, may interfere with or override downstream effects of BAFF signaling. Thus, the stimulation conditions with the largest number of DEGs were the ones with the most differentiated media supernatants at the time of sampling.

Looking at DEGs, we find that long-term stimulation of these B cells with specific cytokines enables strong shifts in the transcriptional landscape, many of which are associated with autoimmunity or B cell malignancies. We see that IL-17 stimulation induces overexpression of *FCRL3*, a known suppressor of BCR activation that is associated with autoimmune disorders, and suppression of *IFI44L*, whose overexpression is associated with autoimmune diseases such as Sjogren’s Syndrome and SLE (Fig 3C)^53,54^. TNFa stimulation induced genes such as *CXCL8* and *POU2AF2*, associated with B cell malignancies, as well as downregulation of NLR genes such as *NLRP4* and *NLRP11*, which work together to dampen inflammation^55–57^. Finally, IL-4 stimulation induces several genes also upregulated at the 4 hour timepoint, such as *CISH* and *CCL17*, but additionally upregulates multiple other genes, such as *IL-17RB*, known to be induced by the combination of IL-4 and CD40L stimulation^58^. High cytokine concentrations for all stimulations except IL-4 and TNFa did not induce any differential gene expression when compared to low concentrations (Supp Fig 3C). Even for IL-4 and TNFa, only a small number of genes were differentially expressed, and so cytokine stimulation concentrations were combined by cytokine (Supp Fig 3D). Cytokine concentration additionally did not impact the amount of IgG secreted into culture supernatant measured via total IgG ELISA for either Bimm pool, and only IL-4 and IL-6 stimulation of Bimm pool 1 samples led to reduced levels of IgG as compared to unstimulated samples (Supp Fig 3E).

As autoimmunity or B cell malignancies are the contexts in which altered glycosylation is often seen, we further probed the transcriptional landscape of these samples to investigate glycogene shifts more specifically^59^. Looking at a PCA of only the panel of 42 known glycogenes, we see that the majority of variance, captured along PC1, describes stimulation by IL-4 (Fig 3D). Additional variance along PC2 describes strong differences between Bimm pools, similarly to observations made at 4 hours of stimulation. These donor differences are similarly strong regardless of stimulation (Supp Fig 3F-K).

To more specifically investigate glycogene profiles induced by stimulation conditions, we turned to a supervised modeling approach called partial least squares discriminant analysis (PLS-DA). We additionally paired PLS-DA classification modeling with feature selection via least absolute shrinkage and selection operator (LASSO), to predict whether a sample has been stimulated or not as compared to an unstimulated condition, using only a reduced set of selected genes from our panel of glycogenes. PLS-DA allows for the utilization and visualization of the variance in the data most tied to the variable of interest, here the stimulation condition. We see that in 10 rounds of 5-fold cross-validation, IL-17, IL-4, and TNFa stimulated conditions separate well from unstimulated controls (Fig 3E). Looking univariately at glycogenes, only IL-4 stimulation induces statistically significantly different glycogene expression (Supp Fig 3L). This suggests that for IL-17 and TNFa stimulation, the unique glycogene profiles are comprised of sub-significant combinatorial changes in glycogene expression, underscoring the interconnected nature of glycogenes, where slight differences in multiple genes in the synthesis pathways may be associated with more noticeable shifts.

Looking at the best models individually, IL-17, IL-4, and TNFa stimulated conditions all exhibit shifts in glycogene expressions whose combinatorial signatures lead to high classification accuracy against unstimulated conditions. Our model is able to separate TNFa-treated cultures from unstimulated cultures, with an accuracy of 0.78 +/− 0.01, with samples across donor pools well-mixed (Fig 3F). Examining the loadings plot of high (|loading value| > 0.02) loadings, where the LASSO-selected features are displayed by contribution to separation and colored by which condition they are enriched in, we see that *ST3GAL2*, *MGAT5*, and *MAN2A2* are the most enriched in TNFa conditions, while *ST3GAL1*, *GNE*, and *FUT8* are enriched in unstimulated conditions, suggesting that galactosylation and fucosylation may differ between conditions (Fig 3G). Selected features highly Spearman correlated with other features are *MAN2B1* and *ST3GAL6*, which are low contributors to the glycogene signature (Supp Fig 3M). IL-17 scores plot shows equally good separation between stimulated and unstimulated samples, with a predictive accuracy of 0.82 +/− 0.04 (Fig 3H). Here, *MGAT5*, *MGAT4B*, and *B4GALT7* are enriched in IL-17 conditions, while *ST3GAL1*, *GNE*, *ST3GAL4*, *ST3GAL1*, and *CMAS* are enriched in unstimulated conditions, among other genes (Fig 3I). Based on these loadings, galactosylation may be higher in IL-17 conditions, while sialylation may be lower. Features highly correlated with selected features include negative correlations between *MGAT5* and *ST3GAL6* as well as *ST3GAL2*, further supporting a lower sialylation hypothesis (Supp Fig 3N). Finally, the IL-4 samples separated best from unstimulated, with a predictive accuracy of 0.98 +/− 0 (Fig 3I). A gene signature of fewer selected genes was necessary for this, suggesting stronger differentiation between samples. IL4 samples were enriched in *B4GALT1* and *ST3GAL1* expression, with unstimulated samples higher in *FUT8*, *B4GALT2,* and *GNE*, indicating that IL-4-treated Bimms may produce IgG with lower levels of fucosylation and galactosylation (Fig 3J). Features enriched in IL-4 stimulated conditions were also highly correlated with sialylation genes such as *ST3GAL4*, suggesting possibly higher levels of sialylation (Supp Fig 3O. These models also perform well against models built with permuted labels or size-matched random features (Supp Fig 3P). Taken together, these results motivated us to examine how transcriptional changes in glycogenes translate into IgG glycosylation.

### Enzyme-linked lectin assay (ELLA) profiling confirms IgG glycosylation changes

Since RNAseq findings at both 4 hours and 14 days of stimulation pointed to altered IgG glycosylation in the Bimm culture, we sought to confirm that these differences translated to the antibody level. Enzyme-linked lectin binding assays (ELLAs) were thus used to measure a variety of glycan moieties in culture supernatants. ELLAs take advantage of the high affinity of lectin proteins for particular glycan moieties, using lectin binding as a proxy for glycan levels^60^. Samples from all high concentration cytokine stimulation conditions at 14 days, as well as low concentration IL-4 and TNFa, both conditions where cytokine concentration induced differential gene expression at 14 days, were run on 8 parallel ELLAs. Lectins were chosen to cover a range of known IgG glycosylation possibilities (Table 3)^61^. Binding profiles across all samples from each lectin primarily showed low correlation, with only MAL-I, PHA-E, and PHA-L having strong significant correlations with each other, indicating that each lectin primarily captured non-overlapping glycan motifs (Supp Fig 4A). Samples from IL-4-stimulated Bimm pool 1 were removed from the analysis, as they had significantly lower total IgG than unstimulated controls via total IgG ELISA, low enough to be nearly undetectable (Supp Fig 3E). Cytokine stimulation showed broad effects on RCA, LCA, and DSL binding across multiple cytokines, and on MAL-I binding for IL-10, IL-6, and TNFa low stimulation (Fig 4A).

**Figure 4.**
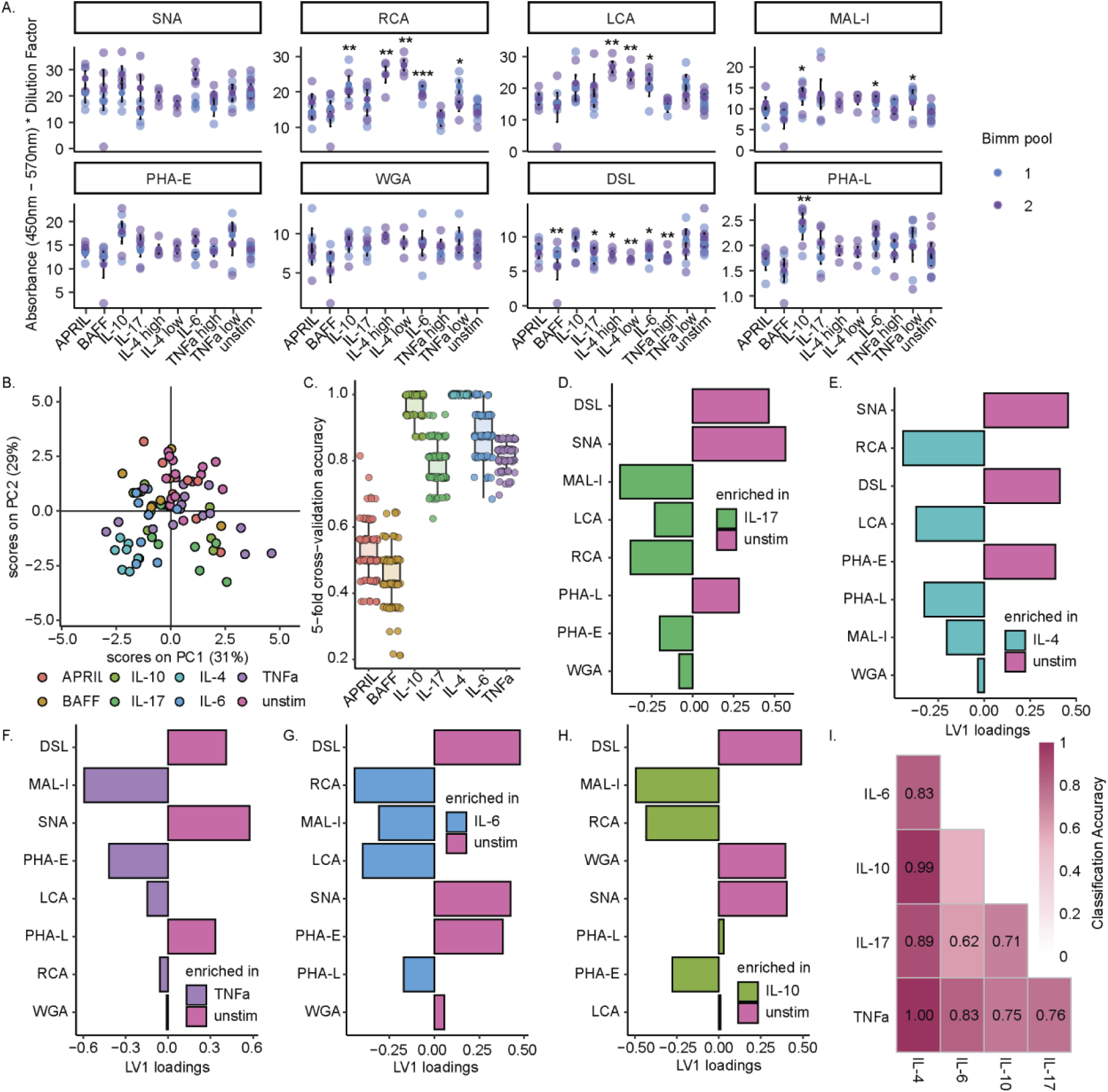
Stimulated conditions exhibit distinct IgG glycosylation profiles. A) Lectin binding levels to IgG from select cytokine stimulation conditions, adjusted for sample dilution. Each dot represents one sample, measured in triplicate, colored by Bimm pool (blue = pool 1, purple = pool 2). Mean values for each Bimm pool are represented by a colored dot, with +/− 1 standard deviation represented by an error bar. Statistical significance between conditions for each lectin was assessed via Kruskal-Wallis test with Benjamini-Hochburg multiple hypothesis corrections. Significant lectins were then assessed pairwise between cytokine stimulation and unstimulated condition for each Bimm pool via Mann-Whitney U test, with Benjamini-Hochberg multiple hypothesis correction (* p < 0.05, ** p < 0.01, *** p < 0.001). B) PCA scores plot for lectin binding across all measured samples. C) 5-fold cross validation accuracy of partial least squares discriminant analysis models built on lectin binding for each pairwise comparison of stimulation to unstimulated condition. Each dot represents the one round of 5-fold cross validation, with boxplots displaying the median value, with lower and upper hinges representing the 25^th^ and 75^th^ percentiles, with the whiskers plotted between the hinges to the largest and smallest values at most 1.5x the interquartile range from their respective hinges. Loadings plots for D) IL17, E) IL4, F) TNFa, G) IL6, H) IL10 stimulation vs unstimulated controls. I) Lower triangle heatmap showing classification accuracy between lectin binding profiles for each cytokine condition combination, colored by accuracy. Values are shown for only models statistically significantly better than models with shuffled labels.

Collectively, these findings reveal a divergent IgG glycosylation profile for certain cytokine-stimulated cultures, with higher LCA binding implying IgG presenting more fucosylation and higher RCA binding implying more galactosylation^61^. While typically increased galactosylation and fucosylation is associated with a less inflammatory profile *in vivo*, studies examining serum IgG glycosylation after seasonal influenza vaccination have found increased galactosylation and fucosylation in the first thirty days post-vaccination^9^. These clonal lines, collected at 14 days post-vaccination, thus might be primed to respond in this way in an inflammatory milieu such as one characterized by the IL-4, IL-6, or TNFa that mimics the increased inflammatory environment in the germinal center post-vaccination.

Alternatively, this divergence from a typical inflammatory glycosylation profile could be due to differences with *in vitro* systems as opposed to *in vivo* settings. Several B cell culture systems have been observed to increase galactosylation when stimulated with pro-inflammatory cytokines such as IFNg and IL-21^19,20^. Meanwhile, *in vivo*, a complex interplay between glycosylation and cytokine milieu takes place, with cytokines impacting glycosylation of secreted IgG, but the binding of differentially-glycosylated IgG also inducing the cellular secretion of various cytokines^62,63^. Notably, all conditions are exposed to the highly inflammatory IL-21 secreted by the feeder cells, so it is possible that an interplay between cytokines is occurring, with cytokines such as IL-6 and IL-4 modulating the effects of IL-21 and leading to slightly increased galactosylation and fucosylation. IL-21 has been shown to, in an *in vitro* B cell system, increase galactosylation substantially more than IL-17, IL-6 or IL-4, so we hypothesize that the effects of IL-21 may prime the cells to raise levels of galactosylation and fucosylation when further stimulated^20^. As IL-10 is the only cytokine considered to be anti-inflammatory within this panel, enhanced RCA binding yet no increase in LCA binding sets it apart from pro-inflammatory stimuli such as IL-4 and IL-6. This binding motif suggests higher levels of galactosylation with no effect on core fucose, which has been associated with an anti-inflammatory profile^64^. While DSL shows differences in binding across multiple cytokine conditions as compared to control, the overall magnitude of DSL binding is low compared to other lectins. Though DSL preferentially binds polyLacNAc, which is not commonly seen on IgG, prior studies examining lectin binding to IgG in the context of *in vitro* cytokine stimulation have seen lower levels of DSL binding to IgG in stimulated as compared to unstimulated conditions^19^.

While various stimulation conditions did induce lectin-binding differences, we checked if stimulations induced distinct profiles overall. While looking within lectins across conditions enables us to infer absolute differences in specific types of glycosylation between conditions, z-scoring across lectins for each sample will enable us to consider compositional differences in overall IgG glycan makeup. A PCA scores plot generated with lectin-binding data z-scored across lectins for each sample revealed a distinct clustering of several conditions separate from unstimulated controls, particularly IL-4, IL-10, and IL-17 (Fig 4B), suggesting there are indeed compositional differences. To identify these lectin fingerprints unique to each condition, we built PLS-DA models on the z-scored lectin data, with the aim of classifying samples as unstimulated or stimulated. Models built on each condition except for APRIL and BAFF stimulation had high (≥ 0.8) mean classification accuracy across 100 rounds of 5-fold cross validation (Fig 4C) and performed well compared to null models (Supp Fig 4B). Multivariate approaches are indeed able to distinguish compositional differences as compared to unstimulated conditions, even for stimulations such as IL-17, which presented few univariate differences in absolute lectin measurements. This underscores the value of considering not only absolute but also relative levels of different types of glycosylation, essentially a glycan signature, when investigating alterations in IgG glycome post-perturbation. Nuanced differences in glycosylation, which we hypothesize could lead to differences in ability of IgG to engage immune cells and stimulate various downstream functional responses, may be missed when considering only absolute univariate differences.

To understand the specific glycan signatures revealed for each stimulation we can examine the PLS-DA model scores and loadings. As supported by high cross-validation scores, all selected PLS-DA models show good separation between stimulated and unstimulated samples (Supp Fig 4C-G), confirming the distinct glycan signatures possessed by each stimulation condition. To understand what those signatures are, we look to the PLS-DA loadings, where higher loading values indicate a more important role for that measurement in the discrimination model.

For all conditions stimulated with classically pro-inflammatory cytokines IL-4, IL-6, IL-17, and TNFa, lectin binding signatures were strongly driven by lower levels of SNA binding and thus sialylation in conjunction with other components of the signature (Fig 4D, E, F, G). This is expected, as a reduction in IgG sialylation has been noted in multiple inflammatory conditions and autoimmune diseases such as rheumatoid arthritis and multiple sclerosis^12,65^. Further, for all pro-inflammatory conditions, LCA binding, implying increased core fucosylation, contributed to separation of stimulated conditions. Notably, the signature of IL-10 stimulation was the only one to show nearly no reliance on LCA binding, suggesting that this anti-inflammatory cytokine did not alter fucosylation (Fig 4H). Other broad signature trends include higher galactosylation, suggested by RCA binding, in all signatures except TNFa stimulation. As low levels of TNFa univariately increased RCA binding (Fig 4A), the lack of RCA binding contribution to this signature must be from the inclusion of both high and low levels of stimulation in the model. TNFa is one of the few cytokines with transcriptional differences between high and low concentration conditions (Supp Fig 3C), with two of the most upregulated genes in the high concentration condition, *CXCL8* and *IL6*, encoding the cytokines IL-8 and IL-6 (Supp Fig 3D). A previous study on IgG glycosylation in Neuromyelitis Optica Spectrum Disorder found a significant negative correlation of both IL-8 and IL-6 levels with IgG galactosylation, suggesting a possible mechanism for RCA binding differences between TNFa stimulation levels^66^.

As several common themes were observed in the signatures derived for each stimulation condition, we built classifier models to determine if conditions were identifiably different from not only unstimulated controls but also from each other. To do so, we built pairwise PLS-DA models between all stimulation conditions, with 10 rounds of 5-fold cross validation. To align with our treatment of different cytokine concentrations in RNAseq analysis, high and low cytokine concentration conditions were combined where available. IL-4 and TNFa treated cultures were clearly distinguishable from all other conditions, while IL-17 and IL-10 were distinguishable from all except IL-6 (Fig 4I). As IL-6 was the least differentiated condition by cytokine Luminex, it stands to reason that it may induce similar glycosylation changes to other conditions (Fig 3B). Thus, despite many of the lectin signatures discriminating stimulated and unstimulated conditions appearing to be similar between stimulations, each stimulation does indeed induce enough of a differentiated signature to be significantly unique, with the exception of IL-6. Finally, as glycogene profiles differed significantly between Bimm pools, we sought to characterize the lectin binding differences between pools. PLS-DA models built to characterize Bimm pool differences found no significant differences in lectin binding profiles between pools 1 and 2, suggesting that observed glycogene differences did not translate directly into IgG glycosylation differences (Supp Fig 4H).

## DISCUSSION

Over time, appreciation for the immunological importance of extra-neutralizing antibody functions such as ADCC, ADNK, and ADCP, among others, has grown. Antibody glycosylation has been recognized as strongly influencing an antibody’s ability to induce these types of functions, and studies have linked alterations in glycosylation and thus in extra-neutralizing antibody functions to numerous immunological perturbations such as vaccination, autoimmune disease, and infection^1,4^. One hypothesis has been that alterations in basal cytokine levels mediate the relationship between immunological perturbations and glycosylation by way of transcriptional regulation^19,67^. However, efforts to study this relationship systematically have been hindered by a variety of issues with model systems, including scarcity of antigen-specific primary B cells and use of non-human systems. Prior studies, focusing on a small number of single cytokine concentrations and a handful of glycogenes, are limited in their ability to uncover complex relationships. Understanding the connection between cytokine stimulation and transcriptional regulation of IgG glycosylation requires an experimentally tractable model system capable of recapitulating antigen-specific human B cell responses to cytokine exposure. Our work characterizes and employs an antigen-specific, immortalized human B cell model to simultaneously characterize IgG glycosylation and glycogene expression in response to cytokine perturbations in a systems-biology framework.

In agreement with prior literature using different model systems, we find that cytokines do indeed alter B cell glycogene expression^19,20^. We expand beyond glycogene investigation to additionally characterize concomitant broad transcriptional profile shifts suggestive of autoimmunity or dysregulated signaling networks, with genes observed to be dysregulated in SLE, Sjogren’s syndrome, and B cell malignancies, among others. Thus, our findings suggest that the glycosylation changes in our model system are occurring in a signaling environment contextually similar to those observed in autoimmune conditions and malignancies. This holds true both for immediate signaling responses, modeled at 4 hours of stimulation, and for longer-term signaling responses where adaptations to chronic cytokine exposure are allowed to occur, around 14 days of stimulation.

Expanding upon literature characterizing differences in RNA levels of glycosyltransferases such as *ST6GAL1*, *NEU1*, and *B4GALT1* after stimulation with IFNg, IL-17, or IL-21, we demonstrate that at both 4 hours and 14 days of cytokine exposure, multivariate modeling approaches enable us to capture unique glycogene signatures of perturbation for an expanded panel of cytokines, in particular IL-4, which to our knowledge has not previously been demonstrated to alter glycogene expression in B cells^19^. *In vitro*, IL-4 affects B cell class-switching, so the glycosylation-related effects of IL-4 expression may further be confounded by immunoglobulin compositional differences^68^. Because Bimm cultures do not class-switch, we are able to directly observe IL-4 effects on glycosylation.

As the glycosylation machinery in a cell is complex, regulated by not only the expression but also the activity and affinity of a series of glycosyltransferases and glycosidases, it becomes important to consider the signaling network as a network in totality and not as individual enzymes. While our system allows for the observation of a small number of significantly differentially expressed glycogenes, a key contribution of this work is the evidence that glycogene signatures, comprised of contributions from multiple genes, are superior in differentiating stimulation conditions from multiple cytokines. Our findings support the idea that lack of statistical significance between conditions when comparing individual glycogenes does not necessarily mean that glycogene expression and thus regulation of glycosylation does not differ between those conditions, rather that the effects may be multivariate. For example, although none of the glycogenes expressed in the IL-17 14 day stimulated cultures were significantly differentially expressed, their contributions to the overall signature allowed us to distinguish treated samples from untreated. Short and long-term responses moreover diverge significantly, additionally confirming the need to consider multiple regulatory levels and timeframes even within the transcriptional space alone.

We find that in glycospace, Bimm pools derived from combinations of clones from different donors exhibit starkly different starting points, but very comparable responses to stimulation. Thus, the Bimm model discussed here is able to capture some of the 50% of glycosylation differences estimated to be heritable^49^. This supports the findings of GWAS studies linking altered IgG glycosylation at a population scale to single nucleotide polymorphisms (SNPs) in a variety of genes encoding for glycosyltransferases but also regulatory transcription factors, regulatory miRNA, and unknown genes^15–17,69^.

We note additionally that combinatorial cytokine stimulation with strong glycogene regulators such as IFNg and IL-4 induces a unique effect in glycogene space, with each stimulation inducing different glycogene up or downregulation, and the combination leading to the additive effect of both cytokines individually. To our knowledge, this is the first *in vitro* evidence of the effects of combinatorial cytokine stimulation on glycosylation. This finding opens future questions as to which other cytokines produce an additive effect, whether pro-inflammatory and regulatory cytokines may lead to effect cancellation, and whether *in vivo* cytokine complexity can be systematically deconstructed in a leave-one-out cytokine stimulation approach to elucidate true drivers of glycosylation differences in complex disease phenotypes.

Since glycogene differences alone are not sufficient to infer IgG glycosylation differences, we leverage parallel ELLAs looking at 8 different types of glycosylation to confirm that transcriptional changes at the 14 day timepoint of stimulation translate into altered IgG glycosylation signatures on the protein level. Five of the cytokines profiled, IL-4, IL-6, IL-17, TNFa, and IL-10, lead to distinct lectin “fingerprints” that agree across pools and distinguish stimulated cultures from unstimulated cultures. The cross-pool agreement suggests shared regulatory responses to these cytokines. As our approach considers glycan profiles compositionally, calculating relative abundance of lectin binding to empaneled lectins within a given sample, a profile high in eg sialylation does not necessarily mean higher absolute sialylation. Rather, it indicates that the major glycosylation type detected in that sample is sialic acid, as opposed to other glycan types measured. IL-10, the sole anti-inflammatory cytokine profiled, exhibits a unique signature with enhanced sialylation but no alterations in fucosylation. The rest of the cytokine panel, primarily pro-inflammatory cytokines, have glycosylation profiles with compositionally less sialylation and more fucosylation. IL-4, a cytokine with complex regulatory functions, that can be anti or pro-inflammatory depending upon context, generates a glycosylation profile similar to more classically pro-inflammatory cytokines such as TNFa.

Unlike in an induced-ASC system, in the Bimm system we do find that IL-4, IL-6, and TNFa affect IgG galactosylation, measured by RCA binding, suggesting key differences in model systems potentially based on priming or immortalization approach. Our findings that pro-inflammatory cytokines induce altered glycosylation align with disease-associated IgG profiles observed *in vivo*. For example, in our prior work we identified altered Fc glycosylation in Lyme disease and tuberculosis, suggesting that cytokine-driven glycosylation changes *in vitro* may reflect the inflammatory environments shaping antibody function during infection^70,71^.

Overall, our findings suggest that Bimms are a tractable model to study cytokine-glycogene-glycosylation links, with the ability to capture glycosylation differences and glycogene differences across multiple glycosylation patterns. This culture system is robust, amenable to long and short-term stimulation, and able to produce transcriptional and proteomic readouts readily, leading to novel findings surrounding IL-4 glycosylation control and combinatorial cytokine stimulation influences on glycogene expression. However, the immortalization strategy may interfere with the ability to observe some types of signaling relationships, here shown by the lack of response to APRIL or BAFF stimulation, which has been observed to alter glycosylation in other culture systems. Additionally, the reliance on IL-21 stimulation in this system does affect the types of glycosylation observable, as IL-21 has been previously observed to alter glycosylation on its own. Interesting future directions with this system would explore the role of cytokine stimulation on individual clones instead of clone pools, further combinatorial cytokine investigations, or the application of complex cytokine milieu to mimic the full serum profile of various autoimmune diseases. Taken together, this work provides a framework for a systematic approach to understanding glycogene regulation of glycosylation, demonstrating links between cytokine-induced glycogene changes and altered IgG glycosylation, and underscoring the importance of considering multivariate glycogene signatures instead of single gene comparisons in complex conditions. This system exhibits the potential to allow for dissection of the mechanisms through which inflammatory environments deriving from the dysregulated signaling in autoimmune or B cell malignancy contexts are able to tune antibody effector functions by way of glycosylation shifts.

## Author Contributions

Conceptualization and Methodology: C.D.W., D.A.L.; Resources: T.A.W., K.L.B., R.A.K.; Investigation: C.D.W., M.S.A., C.W.; Data Curation: C.D.W., C.W.; Formal Analysis: C.D.W., M.S.A., D.A.L.; Visualization: C.D.W.; Writing – Original Draft: C.D.W.; Writing – Review & Editing: C.D.W., M.S.A., C.W., K.L.B., D.A.L.; Supervision: D.A.L.

## Acknowledgements

The authors thank Glenn Paradis, Michele Griffin, and Michael Tramontanis of the Koch Institute Flow Cytometry Core for their technical expertise and support, as well as MIT’s BioMicroCenter Core, particularly Dr. Stuart Levine and Avanyish Toniappa. The authors also thank Julia Zhong, Dr. Diana Gong, Dr. Anisha Datta, Dr. Krista Pullen, Dr. Laura Bahlmann, and Dr. Brian Joughin for their technical expertise and constructive discussions. We gratefully acknowledge the gift of the feeder cell line YK6-CD40Lg-IL21 from Dr. Daniel Hodson and Miriam Di Re, Hodson Lab, Wellcome-MRC Cambridge Stem Cell Institute, University of Cambridge.

This research was supported in part by the Intramural Research Program of the National Institutes of Health (NIH). The contributions of the NIH author(s) are considered Works of the United States Government. The findings and conclusions presented in this paper are those of the author(s) and do not necessarily reflect the views of the NIH or the U.S. Department of Health and Human Services. This project has been funded in in part with federal funds from the National Institute of Allergy and Infectious Diseases, the NIH, and the Department of Health and Human Services under contract No. 75N93019C00071, grant No. AI181898, and grant No. U19AI135995. We would also like to thank the Fairbairn Family Fund for supporting this work, as well as Army ICB UARC Contract W911NF-19-D-0001. Funding was also provided by the National Cancer Institute of the NIH under award P30-CA14051, and this work was further supported by a graduate student fellowship from the MIT-Takeda Fellow Program. Several experimental schematics were created with BioRender.

## Declaration of Interests

The authors declare that the research was conducted in the absence of any commercial or financial relationships that could be construed as a potential conflict of interest.

## METHODS

### Cell Culture

The YK6-CD40Lg-IL21 feeder line “YK6” was cultured in Advanced Roswell Park Memorial Institute (Advanced RPMI 1640, ThermoFisher) medium containing 10% fetal bovine serum (FBS, ThermoFisher) and 1% penicillin/streptomycin (ThermoFisher), “YK6 media”. YK6 cells were irradiated with 30 Gy via Gammacell irradiator prior to banking at a density of 4e6 viable cells/mL (vc/mL) in freezing media (45 mL FBS, 5 mL dimethyl sulfoxide, Sigma Aldrich).

Influenza H7 HA antigen-specific immortalized B cell (Bimm) lines F2-2, F2-11, F2-21, F5-16, F5-17, F5-24 were cultured in Iscove’s Modified Dulbecco’s Medium (IMDM, ThermoFisher) supplemented with 10% FBS and 1% penicillin/streptomycin. Cells were cultured on irradiated YK6 cells (2e6 vc/T75 flask or 200,000 vc/well in 12 well plates) at an initial density of 100,000-200,000 cells/well. Cell counting was done via hemocytometer on an EVOS M5000 imaging system using the green fluorescent channel to count only Bimm cells. Culture media was changed every 3 to 4 days, with new irradiated YK6 cells added every 7 days. After 14 days of culture, Bimms were moved to fresh flasks or plates containing new irradiated YK6 cells. To transfer cells, the surface of the culture was rinsed vigorously with fresh media to remove the Bimms while disturbing as little of the adherent YK6 layer as possible, and the resulting culture volume was transferred to the new flask or plate.

All cell lines used in this study were maintained in a humidified incubator (5% CO2) at 37C. All cell lines used in this study were confirmed free of mycoplasma bacteria via MycoAlert mycoplasma detection kit (Lonza).

### IFNg and BCR stimulation experiments

One clone, F2-21, was seeded at 220,000 vc/mL in 12 well plates coated with YK6 feeder cells. Cells were allowed to grow for 9 total days, with YK6 feeder supplement at day 6. At 48, 24, or 4 hours prior to experimental end, media was changed to fresh media supplemented with 50 ng/mL IFNg (PeproTech), 5 ng/mL a-IgG (anti-IgG F(ab’)2, Invitrogen) for BCR stimulation, both, or neither (n = 4 replicates per timepoint and condition). Bimms were isolated via media rinsing followed by CD19 positive selection and immediately lysed for RNAseq.

### Bimm immunoglobulin secretion

Bimms were pooled by donor at equal concentrations and each Bimm pool was seeded at 100,000 vc/mL in 12 well plates coated with YK6 feeder cells. 3 replicates per pool were seeded for each of the 7 desired timepoints. Cells were allowed to grow for 10 days, with a YK6 feeder supplementation at day 7. On day 10, media was changed for all wells. Supernatant was collected from new wells at 4, 6, 8, 12, 18, 24, and 48 hours post media change, with any given well harvested once. Supernatant was assayed for presence of immunoglobulin using a human total IgG ELISA kit (Invitrogen/Thermo Fisher Scientific), according to manufacturer’s instructions.

### Bimm stimulation experiments

200,000 irradiated YK6 feeder cells were seeded per well on 12 well plates in YK6 media and allowed to adhere for 24 hours. Bimm lines were then pooled by donor at equal concentrations and each Bimm pool was seeded at 100,000 vc/mL in the 12 well plates pre-coated with feeder cells. Bimm pools were grown for 2 days (14 day stimulation) or 16 days (4 hour stimulation). Media was changed at the noted time to media containing IL-4, IL-6, IL-10, IL-17, TNF-a, IFN-g, APRIL, and BAFF (PeproTech) at both high and low concentrations (Table 1) for 14 days or 4 hours. For stimulation experiments, the FBS supplement in the media was changed to Ultra low IgG FBS (Invitrogen), to mitigate the effects of exogenous IgG. For cultures stimulated to engage the B cell receptor (BCR), anti-IgG F(ab’)2 (5 mg/mL, Invitrogen) was added to the media as well for the first 3 days of stimulation (14 day stimulation) or the entire 4 hours (4 hour stimulation). 100,000 irradiated YK6 cells were added to each well after 7 days of YK6 culture, and all cultures were moved to new irradiated YK6 12-well plates at day 14 of YK6 culture. Cell culture supernatants at the end of experiment day 16 were collected and stored at −80C. Cells were immediately isolated and purified as described below and submitted for RNA-sequencing. 5 experimental replicates were run, with each replicate containing one instance of each condition. Replicates were seeded on separate days from the same Bimm cultures, with all steps including Bimm pooling performed independently per replicate. For the 14 day stimulation experiment, a 6^th^ replicate was run to ensure all conditions were represented in the final data at least three times. At experimental end, supernatant was collected for downstream analyses and cells were isolated via the Bimm purification protocol.

### Bimm purification

To purify Bimms from co-culture with irradiated YK6 cells, CD19 positive selection was performed via EasySep Human CD19 Positive Selection kits (STEMCELL Technologies). Supernatant was removed from all wells of 12-well plates, and 500 uL EasySep buffer (ion-free PBS, 2% heat-inactivated FBS, 1 mM EDTA) was added to each well and used to vigorously rinse the well surface. Cultures were transferred into a 96 well plate and resuspended in 50 uL EasySep buffer. Positive selection was performed in 96 well plates according to manufacturer instructions using the EasyPlate magnet. Cells were then resuspended in 300 uL RNAse-free water per well and transferred into a 96 well deep-well plate. 300 uL lysis buffer was added per well and thoroughly mixed. 25 uL B-mercaptoethanol (Sigma Aldrich) was added as well, and samples were immediately transferred to the MIT BioMicroCenter sequencing core facility for further analysis.

### RNA-sequencing

RNA-sequencing was performed by the MIT BioMicroCenter sequencing core facility. RNA was extracted using the Chemagic360 (Perkin Elmer). mRNA was then extracted via PolyA isolation (New England BioLabs) and sequenced with the Element AVITI (Element Biosciences) using 150 nt flowcells. RNA from each replicate experiment was extracted on different days, immediately following cell lysis, while multiple experiments were run together per flowcell. Sequences from different flow cell lanes for each end of the paired-end sequencing results were merged. Read quality was checked using fastqc (v 0.11.5) and sequences with low per-base sequence quality (below 28) were trimmed using fastp (v 0.20.0) with default settings. Quality was again checked for trimmed reads using fastqc. Sequences were aligned to the GRCh38.p14 reference genome, indexed using bowtie2 (v 2.3.5.1) via tophat (v 2.1.1). Samtools (v 1.10) was used to index accepted hits for only samples with >70% alignment, and HTSeq was used to generate count data. Mitochondrial genes were filtered out, as were unassigned genes and genes with a read count less than 10.

### Flow cytometry

Irradiated YK6 cells were stained with CellTrace Violet (ThermoFisher) according to the manufacturer’s instructions. Stained irradiated YK6 cells were then plated at a density of 200,000 vc/mL in 1 mL media per well for 12 well plates and left to adhere overnight. For assessment of the isolation protocol, Bimms were isolated according to the protocol above. As a baseline comparison, co-cultures of Bimms and irradiated YK6 cells were lifted from 12-well plates using 0.25% Trypsin-EDTA (Fisher Scientific). Cell cultures of interest were resuspended in FACS buffer (PBS / 2% FBS) and blocked for 10 minutes at room temperature with Human TruStain FcX (BioLegend Inc.). Cells were then resuspended in FACS buffer and stained with Zombie Near-IR (ThermoFisher) for viability according to manufacturer instructions. Single-stain controls were additionally generated. All samples were fixed in 4% paraformaldehyde for 30 minutes on ice before resuspension in FACs buffer and transfer to flow cytometry tubes. Data was immediately acquired on a FACS LSR Fortessa HTS-2 at the Koch Institute Flow Cytometry Core. Compensation and gating were performed in FlowJo (10.8.2).

### Antibody quantification via ELISA

Concentrations of total IgG in the supernatant were determined by human total IgG ELISA (Invitrogen) according to manufacturer instructions. Supernatant was diluted 1:5, 1:10, or 1:20 to fall within a linear dilution range and within the range of the manufacturer-provided standard curve.

### Cytokine quantification via Luminex

Media supernatants from 3 experimental replicates were assayed with custom Luminex kits (R&D Systems) comprised of detection beads for APRIL/TNFSF13, BAFF/BlyS/TNFSF13B, GM-CSF, IFN-gamma, IL-4, IL-6, IL-10, IL-17/IL-17A, and TNF-alpha. Volumes were adjusted to 12.5 uL of sample and reagents per well to adapt kits to a 384-well format. Samples were diluted 1:2, 1:10, and 1:20 in diluent and run in technical duplicate on a Bio-Plex 3D Suspension Array System (Bio-Rad). Data were adjusted to account for background in R by subtracting the median fluorescent intensity (MFI) of a media sample, IMDM supplemented with 10% Ultra low IgG FBS and 1% penicillin/streptomycin, from the mean MFI of each sample’s technical replicates. All negative values were adjusted to one, and data were then log10 transformed.

### Enzyme-linked lectin assays (ELLA)

Media supernatants from 3 experimental replicates were assayed with a custom enzyme-linked lectin assay (ELLA). 384-well plates were coated with 4 ug/mL Protein A (ProA, ThermoFisher) and incubated for 4 hours shaking at room temperature., except for wells designated as standard wells. Protein A binding allows for the isolation of immunoglobulins from cell culture supernatant, as other glycosylated supernatant components such as cytokines would otherwise interfere with ELLA readouts. After 3 incubation hours, bovine fetuin B (Sigma Aldrich) at a concentration of 2.5 ug/mL in PBS was prepared and serially diluted 1:2 to generate a nine-point standard curve, with the tenth point being plain PBS. This standard curve was added to the standard wells, and the plate was left to finish the final hour of incubation. The plate was then washed three times using a wash buffer of PBS and 0.2% Tween20 (PBST). The plate was blocked for 1 hour shaking at room temperature with SynBlock blocking buffer (BIO-RAD) before another three washes in PBST. Media supernatant diluted 1:25 or 1:5 in PBS, based on sample titer, was added to ProA wells in triplicate, and PBS was added to standard wells. The plate was then incubated overnight shaking at 4C. After 3 PBST washes, a solution of lectin (Vector Labs) in binding buffer (DPBS supplemented with 10 mM HEPES) was added to each well and the plate incubated for 1 hour shaking at room temperature. Lectin concentrations, sources, and specificities are noted in Table 3. After 3 PBST washes, Streptavidin-HRP (Strep B, R&D Systems) was added to all wells and the plate incubated for 30 minutes shaking at room temperature. After 3 PBST washes, ELISA TMB substrate (R&D Systems) was added to the plate and allowed to incubate until appropriate standard curve signal arose, 5-20 minutes. 2N sulfuric acid (R&D Systems) was then added to stop the reaction, and the plate was read on via plate reader at 450 nm and 570 nm. Standards and samples with 570 nm readings over 0.15 were discarded, as occasionally precipitates formed due to the differences in kinetics between standard color development and sample color development. Only samples that fell inside a linear range of standard readings were considered for analysis.

### Univariate Analyses

All univariate plots were visualized and analyzed using R (version 4.3.1). Univariate plots show the dilution-adjusted absorbance values for ELLA assays, visualized as dot plots, with solid dots denoting the mean value per group and whiskers denoting +/− one standard deviation. Unless noted, statistical significance between measurements was calculated with the Mann-Whitney nonparametric test, with p-values then corrected for multiple hypothesis testing via Benjamini-Hochburg. Heatmaps were generated via ‘pheatmap’ from ‘pheatmap’ (v 1.0.12). Figures were edited only for page positioning, legend color, and size in Adobe Illustrator (v2020) for clearer visualization.

### RNAseq Data Analysis

Immunoglobulin transcripts were removed from the counts matrix prior to analysis. Differential gene expression was performed using DESeq2 (v 1.40.2), with a design matrix employed to account for Bimm pool, sequencing plate, cytokine, and BCR stimulation. Differentially expressed genes (DEGs) were determined via contrasts of interest, with log2 fold changes shrunk using lfcShrink and the adaptive shrinkage estimator ‘ashr’. DEGs were chosen as those with an adjusted p-value < 0.05 and an absolute log2 fold change greater than 1 for all analyses at 4-24 hours, or greater than 0.58 at 14 days, unless otherwise noted. For visualization purposes, variance stabilizing transformation was utilized and sequencing plate effects were regressed out using limma’s ‘removeBatchEffect’ function (v 3.58.1). When applicable, transcription factor activity was inferred from normalized log2 gene counts using ‘run_ulm’ from the R package ‘decoupleR’ (v 2.6.0), paired with regulatory networks from the ‘CollecTri’ resource. DecoupleR allows for the inference of transcription factor activity from transcriptomics data based on signed regulatory knowledge of transcription factor control of gene expression. Gene set enrichment analysis (GSEA) was performed using ‘fgsea’ from R package ‘fgsea’ (v 1.26.0) on genes ranked by log2 fold change, with pathways from the Reactome resource (v 1.84.0). Pathways were considered significant with an adjusted p-value < 0.05.

### Multivariate Analyses

Multivariate analyses included both lectin and RNAseq measurements. For unsupervised principal component analysis (PCA) models, data were z-scored by gene for RNAseq data or by sample for lectin data. PCA was performed using ‘prcomp’ from ‘stats’ (v 4.3.1) and visualized via ggplot (v 3.5.1). For classification models, partial least squares discriminant analysis (PLS-DA) was run using ‘train_ropls’ and visualized using ‘visualize_ropls_scores’ and ‘visualize_ropls_loadings_bar’ from the systemsseRology R package (v1.1). For PLS-DA on glycogenes, least absolute shrinkage and selection operator (LASSO) regularization was performed first for feature reduction using the ‘systemsseRology’ function ‘select_lasso’ to run feature selection 100 times. Features selected at least 80 of the 100 rounds were retained for model building to reduce overfitting risk.

Generalizability of PLS-DA models was assessed in a 10 or 100 round five-fold cross validation framework as noted, where each round consists of 100 repeats of feature selection per fold. The mean of these rounds was compared to null models built by permuting group labels or selecting random, size-matched features. To generate these null models, 10 or 100 rounds of 5-fold cross validation with 100 permutations and 100 random feature trials per round were performed, unless otherwise noted, still with 100 repeats of feature selection per fold, and p-values were calculated using Mann-Whitney U tests and multiple-hypothesis corrected via Benjamini-Hochburg.

### Correlation Networks

To account for features highly correlated with selected features in the RNAseq PLS-DA models, features with high absolute Spearman correlations (Spearman R > |0.7|, p < 0.05) to selected features were retained. Correlations were calculated with the function ‘correlate’ in ‘corrr’ (v 0.4.4), with p-values calculated via ‘cor.test’ and multiple hypothesis corrected via Benjamini-Hochburg. ‘ggraph’ (v 2.2.1) and ‘igraph’ (v 2.1.2) were used to visualize correlation networks. Node coloring and positioning was adjusted in Adobe Illustrator (v2020) for improved visualization only. Edge color represents the Spearman R correlation strength, while node color represents feature status, with white nodes as co-correlate features and colored nodes as LASSO-selected features.

## Notes

### Competing Interest Statement

The authors have declared no competing interest.

