## Supplemental Figures for "Multivariate analysis of glycogenes reveals coordinated regulation of immunoglobulin glycosylation in an immortalized human B cell system"

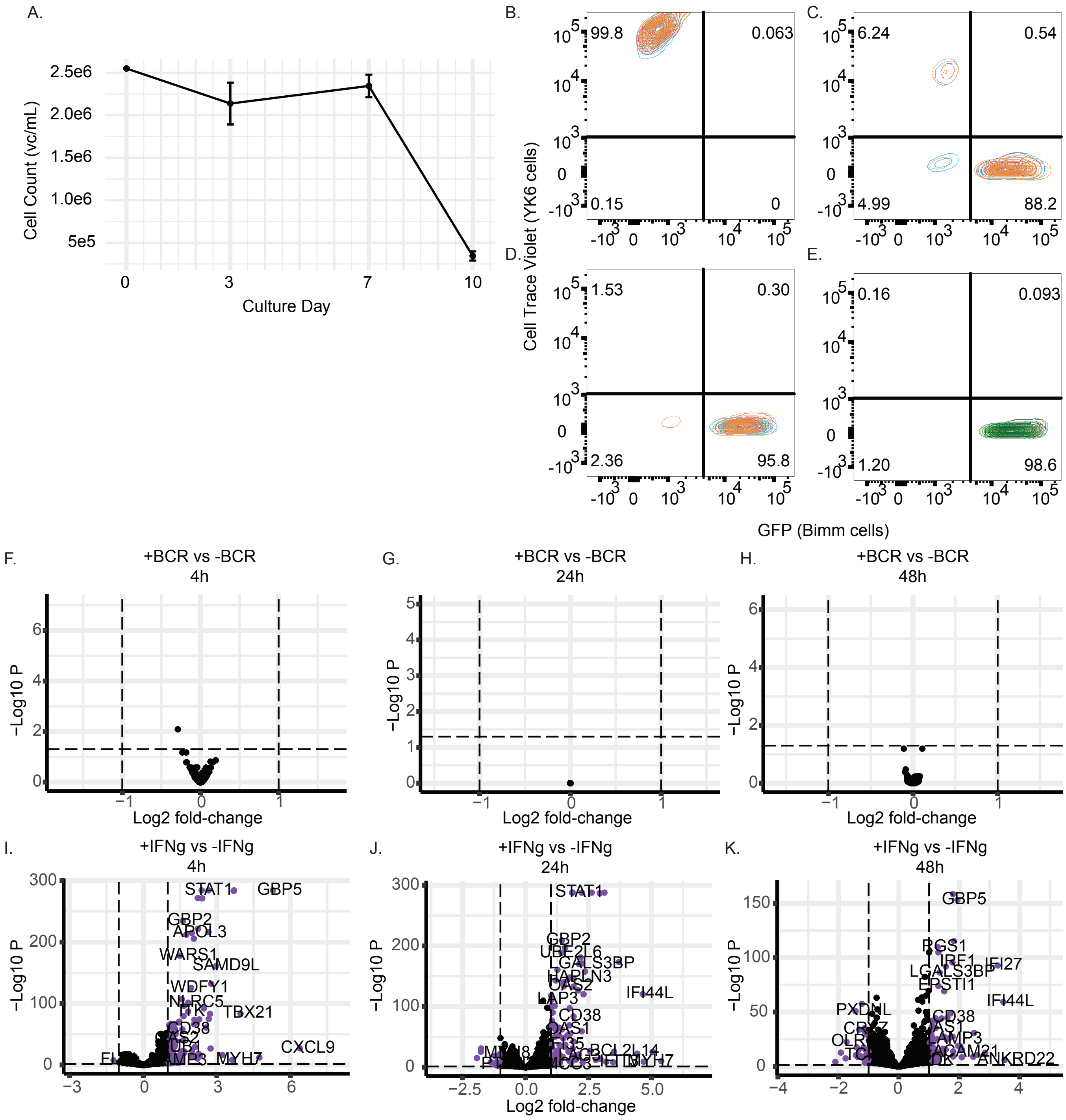
A.
Cell Count (vc/mL)
Culture Day
B.
C.
D.
E.
Cell Trace Violet (YK6 cells)
GFP (Bimm cells)
99.8
0.063
0.15
0
6.24
0.54
4.99
88.2
1.53
0.30
2.36
95.8
0.16
0.093
1.20
98.6
F.
+BCR vs -BCR
4h
G.
+BCR vs -BCR
24h
H.
+BCR vs -BCR
48h
−Log10 P
Log2 fold-change
I.
+IFNg vs -IFNg
4h
J.
+IFNg vs -IFNg
24h
K.
+IFNg vs -IFNg
48h
STAT1
GBP5
GBP2
APOL3
WARS1
SAMD9L
WDFY1
NLRC5
TBX21
CD38
CXCL9
MYH7
GBP2
LGALS3BP
IFI44L
CD38
OAS1
BCL2L14
MYH7
GBP5
IFI27
LGALS3BP
EPSTI1
IFI44L
CD38
LAMP3
ANKRD22
PXDNL

**Supplemental Figure 1.** A) Growth curve for irradiated YK6 cells, n = 3 replicates. Mean +/- 1 standard deviation shown. Flow cytometry contour plots for B) irradiated YK6 cells (n = 3), C) trypsin-lifted co-culture (n = 3), D) media-rinsed co-culture (n = 3), or E) CD19 positively selected Bimms (n = 5). Differentially expressed genes for a BCR stimulated condition vs unstimulated at F) 4h, G) 24h, and H) 48h of stimulation, adjusted $p < 0.05$, $|\log2 \text{ fold-change}| > 1$. Differentially expressed genes for an IFNg stimulated condition vs unstimulated at I) 4h, J) 24h, and K) 48h of stimulation, adjusted $p < 0.05$, $|\log2 \text{ fold-change}| > 1$.

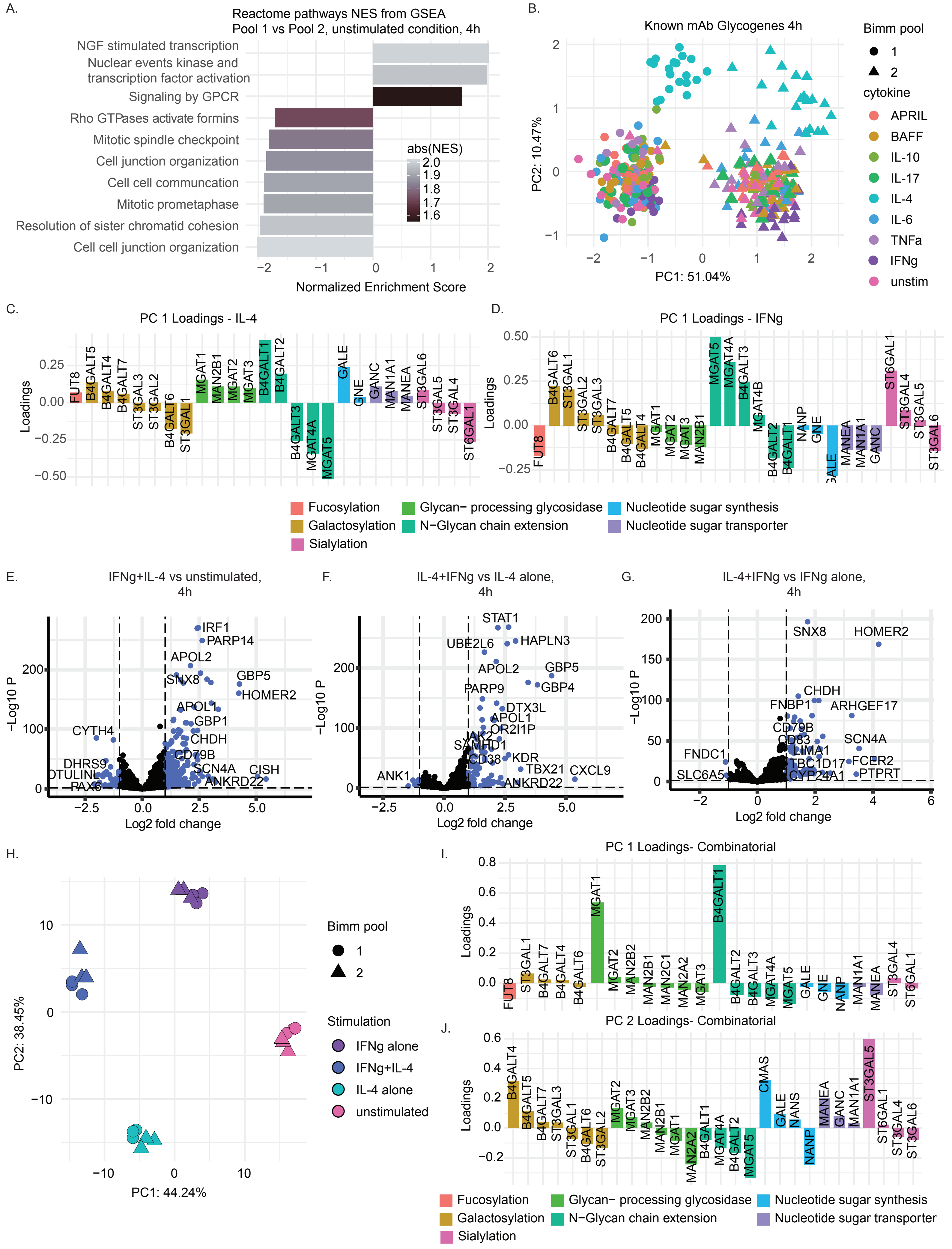

A.
Reactome pathways NES from GSEA
Pool 1 vs Pool 2, unstimulated condition, 4h
NGF stimulated transcription
Nuclear events kinase and transcription factor activation
Signaling by GPCR
Rho GTPases activate formins
Mitotic spindle checkpoint
Cell junction organization
Cell cell communcation
Mitotic prometaphase
Resolution of sister chromatid cohesion
Cell cell junction organization
abs(NES)
2.0
1.9
1.8
1.7
1.6
−2
−1
0
1
2
Normalized Enrichment Score
B.
Known mAb Glycogenes 4h
PC2: 10.47%
PC1: 51.04%
Bimm pool
1
2
cytokine
APRIL
BAFF
IL-10
IL-17
IL-4
IL-6
TNFa
IFNg
unstim
C.
PC 1 Loadings - IL-4
Loadings
FUT8
B4GALT5
B4GALT4
B4GALT7
ST3GAL3
ST3GAL2
B4GALT6
ST3GAL1
MGAT1
MAN2B1
MGAT2
MGAT3
B4GALT1
B4GALT2
B4GALT3
MGAT4A
MGAT5
GALE
GNE
GANC
MAN1A1
MANEA
ST3GAL6
ST3GAL5
ST3GAL4
ST6GAL1
D.
PC 1 Loadings - IFNg
Loadings
FUT8
B4GALT6
ST3GAL1
ST3GAL2
ST3GAL3
B4GALT7
B4GALT5
B4GALT4
MGAT1
MGAT2
MGAT3
MAN2B1
MGAT5
MGAT4A
B4GALT3
MGAT4B
B4GALT2
B4GALT1
NANP
GNE
GALE
MANEA
MAN1A1
GANC
ST6GAL1
ST3GAL4
ST3GAL5
ST3GAL6
Fucosylation
Galactosylation
Sialylation
Glycan− processing glycosidase
N−Glycan chain extension
Nucleotide sugar synthesis
Nucleotide sugar transporter
E.
IFNg+IL-4 vs unstimulated, 4h
−Log10 P
Log2 fold change
IRF1
PARP14
APOL2
SNX8
GBP5
HOMER2
APOL1
GBP1
CHDH
CD79B
SCN4A
CISH
ANKRD22
CYTH4
DHRS9
OTULINL
PAX6
F.
IL-4+IFNg vs IL-4 alone, 4h
−Log10 P
Log2 fold change
STAT1
UBE2L6
HAPLN3
APOL2
GBP5
GBP4
PARP9
DTX3L
APOL1
OR2I1P
JAK2
SAMHD1
CD38
KDR
TBX21
CXCL9
ANKRD22
ANK1
G.
IL-4+IFNg vs IFNg alone, 4h
−Log10 P
Log2 fold change
SNX8
HOMER2
CHDH
FNBP1
ARHGEF17
CD79B
CD83
SCN4A
LIMA1
FNDC1
TBC1D17
FCER2
SLC6A5
CYP24A1
PTPRT
H.
PC2: 38.45%
PC1: 44.24%
Bimm pool
1
2
Stimulation
IFNg alone
IFNg+IL-4
IL-4 alone
unstimulated
I.
PC 1 Loadings- Combinatorial
Loadings
FUT8
ST3GAL1
B4GALT7
B4GALT4
B4GALT6
MGAT1
MGAT2
MAN2B2
MAN2B1
MAN2C1
MAN2A2
MGAT3
B4GALT1
B4GALT2
B4GALT3
MGAT4A
MGAT5
GALE
GNE
NANP
MAN1A1
MANEA
ST3GAL4
ST6GAL1
J.
PC 2 Loadings- Combinatorial
Loadings
B4GALT4
B4GALT5
B4GALT7
ST3GAL3
ST3GAL1
B4GALT6
ST3GAL2
MGAT2
MGAT3
MAN2B2
MAN2B1
MGAT1
MAN2A2
B4GALT1
MGAT4A
B4GALT2
MGAT5
CMAS
GALE
NANS
NANP
MANEA
GANC
MAN1A1
ST3GAL5
ST6GAL1
ST3GAL4
ST3GAL6
Fucosylation
Galactosylation
Sialylation
Glycan− processing glycosidase
N−Glycan chain extension
Nucleotide sugar synthesis
Nucleotide sugar transporter

**Supplemental Figure 2.** A) Top 10 Reactome gene set enriched pathways for unstimulated Bimm pool 1 vs Bimm pool 2 samples at 4h post media-change. B) PCA scores plot of known glycogenes at 4h of stimulation. Principal components loadings for PC1 for C) IL-4 and D) IFNg PCAs, with loadings colored by the glycogene family they belong to. Differentially expressed genes for a combinatorial IFNg+IL4 stimulated condition vs E) unstimulated, F) IL4 stimulated, or G) IFNg stimulated, adjusted $p < 0.05$, $|\log2 \text{ fold-change}| > 1$. H) PCA scores plot of all differentially expressed genes identified between IFNg+IL4 stimulated samples and either unstimulated, IL4 only stimulated, or IFNg only stimulated samples. Principal components loadings for I) PC1 and J) PC2 for combinatorial stimulation experiment, corresponding to scores plot in Figure 2. Loadings are colored by the glycogene family they belong to. For all loadings plots, only genes with $|\text{loading value}| > 0.02$ are shown.

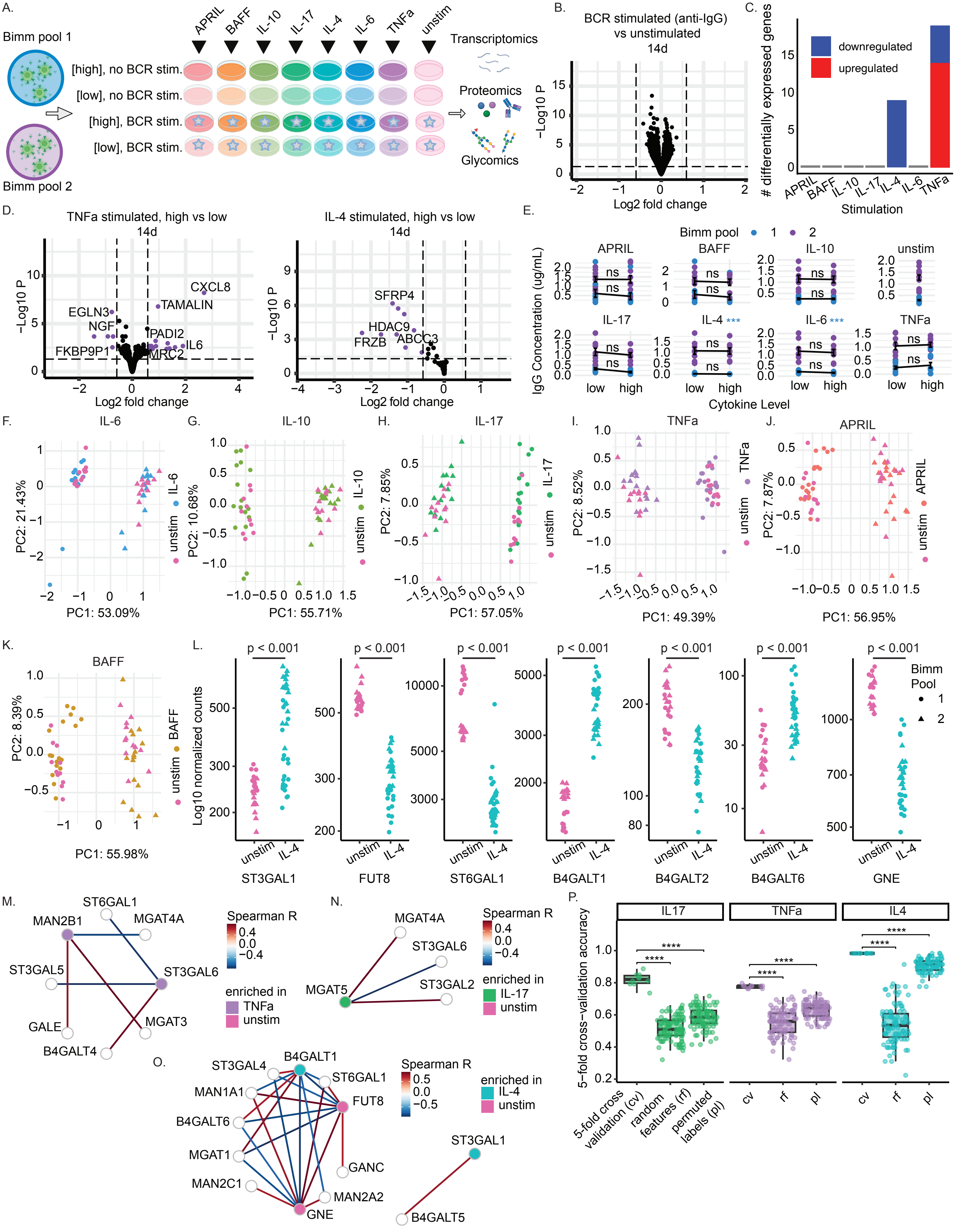

A.
Bimm pool 1
Bimm pool 2
APRIL
BAFF
IL-10
IL-17
IL-4
IL-6
TNFa
unstim
[high], no BCR stim.
[low], no BCR stim.
[high], BCR stim.
[low], BCR stim.
Transcriptomics
Proteomics
Glycomics
B.
BCR stimulated (anti-IgG) vs unstimulated
14d
−Log10 P
Log2 fold change
C.
### differentially expressed genes
downregulated
upregulated
Stimulation
D.
TNFa stimulated, high vs low
14d
CXCL8
TAMALIN
EGLN3
NGF
PADI2
IL6
FKBP9P1
MRC2
IL-4 stimulated, high vs low
14d
SFRP4
HDAC9
FRZB
ABCC3
E.
Bimm pool
1
2
IgG Concentration (ug/mL)
ns
***
low
high
Cytokine Level
F.
G.
H.
I.
J.
PC1: 53.09%
PC2: 21.43%
PC1: 55.71%
PC2: 10.68%
PC1: 57.05%
PC2: 7.85%
PC1: 49.39%
PC2: 8.52%
PC1: 56.95%
PC2: 7.87%
K.
PC1: 55.98%
PC2: 8.39%
L.
p < 0.001
Log10 normalized counts
ST3GAL1
FUT8
ST6GAL1
B4GALT1
B4GALT2
B4GALT6
GNE
Bimm Pool
M.
ST6GAL1
MAN2B1
MGAT4A
ST3GAL5
ST3GAL6
GALE
MGAT3
B4GALT4
Spearman R
enriched in
TNFa
unstim
N.
MGAT4A
ST3GAL6
MGAT5
ST3GAL2
IL-17
O.
ST3GAL4
B4GALT1
ST6GAL1
MAN1A1
FUT8
B4GALT6
MGAT1
GANC
MAN2C1
MAN2A2
GNE
ST3GAL1
B4GALT5
IL-4
P.
IL17
TNFa
IL4
5-fold cross−validation accuracy
****
5-fold cross validation (cv)
random features (rf)
permuted labels (pl)
cv
rf
pl

**Supplemental Figure 3.** A) Schematic of stimulation experiment, with 7 cytokines supplemented at either a high or low concentration, both with and without BCR stimulation, for two Bimm pools. B) Differentially expressed genes for a BCR stimulated condition vs unstimulated, at 14d, adjusted $p < 0.05$, $|\log 2 \text{ fold-change}| > 0.58$. C) Number of differentially expressed genes per stimulation, colored by whether genes were upregulated or downregulated by a high concentration of that stimulation as compared to a low concentration, adjusted $p < 0.05$, $|\log 2 \text{ fold-change}| > 0.58$. D) Differentially expressed genes for TNFa and IL4 high concentration conditions compared to low concentration conditions, adjusted $p < 0.05$, $|\log 2 \text{ fold-change}| > 0.58$. E) Media concentration of total IgG measured by total human IgG ELISA over time per sample, samples measured in triplicate. Samples plotted by cytokine stimulation condition and level. Statistical significance between cytokine level for each cytokine and Bimm pool combination was assessed via Mann-Whitney U test, with Benjamini-Hochberg multiple hypothesis correction. Statistical significance between cytokine stimulation and unstimulated condition for each Bimm pool was assessed via Mann-Whitney U test, with Benjamini-Hochberg multiple hypothesis correction (*** $p < 0.001$). PCA scores plots built on known glycogenes for stimulated vs unstimulated samples at 14d, for F) IL6, G) IL10, H) IL17, I) TNFa, J) APRIL, and K) BAFF stimulation. For all scores plots, score shape corresponds to Bimm pool (circle = Bimm pool 1, triangle = Bimm pool 2). L) Log10 normalized counts for glycogenes statistically significantly different from unstimulated controls in any stimulated conditions. Statistical significance determined via DESeq2. Spearman correlations for LASSO-selected genes for M) TNFa PLS-DA, N) IL-17 PLS-DA, and O) IL-4 PLS-DA, with nodes colored by which condition they were enriched in, and edges colored by Spearman R. P) 5-fold cross validation accuracy of partial least squares discriminant analysis models built on known glycogenes for each pairwise comparison of IL17, TNFa, and IL4 stimulated to unstimulated conditions. Each dot represents the one round of 5-fold cross validation, with boxplots displaying the median value, with lower and upper hinges representing the 25$^{th}$ and 75$^{th}$ percentiles, with the whiskers plotted between the hinges to the largest and smallest values at most 1.5x the interquartile range from their respective hinges. Accuracy metrics shown for each stimulation are mean 5-fold cross-validation accuracy, accuracy of null models generated with size-matched random features, or accuracy of null models generated with permuted group labels. 10 trials of 5-fold cross validation were run, with 100 trials of permutation or random features per trial. Statistical significance was determined via pairwise Mann-Whitney U tests, (**** $p < 0.0001$).

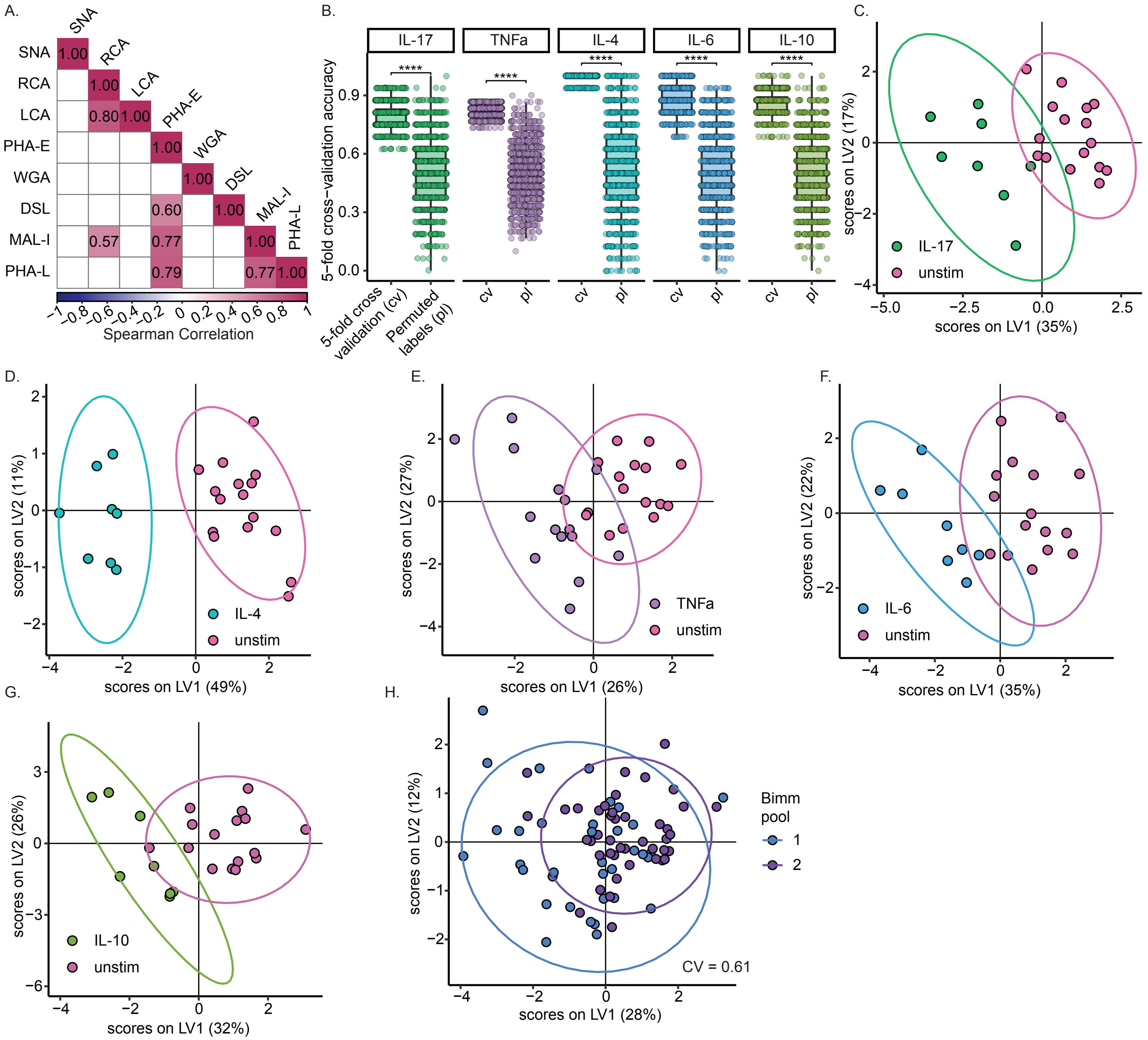
A.
SNA
RCA
LCA
PHA-E
WGA
DSL
MAL-I
PHA-L
1.00
0.80
0.60
0.57
0.77
0.79
0.77
Spearman Correlation
B.
IL-17
TNFa
IL-4
IL-6
IL-10
****
5−fold cross−validation accuracy
5-fold cross validation (cv)
Permuted labels (pl)
cv
pl
C.
IL-17
unstim
scores on LV1 (35%)
scores on LV2 (17%)
D.
IL-4
unstim
scores on LV1 (49%)
scores on LV2 (11%)
E.
TNFa
unstim
scores on LV1 (26%)
scores on LV2 (27%)
F.
IL-6
unstim
scores on LV1 (35%)
scores on LV2 (22%)
G.
IL-10
unstim
scores on LV1 (32%)
scores on LV2 (26%)
H.
Bimm pool
1
2
CV = 0.61
scores on LV1 (28%)
scores on LV2 (12%)

**Supplemental Figure 4.** A) Lower triangle heatmap of Spearman correlations between lectin binding for each lectin across all stimulation conditions, colored by Spearman R. Correlation values are shown for only statistically significant correlations. B) 5-fold cross validation accuracy of partial least squares discriminant analysis models built on lectin binding data for each pairwise comparison of IL17, TNFa, IL4, IL6, and IL10 stimulated to unstimulated conditions. Each dot represents the one round of 5-fold cross validation, with boxplots displaying the median value, with lower and upper hinges representing the $25^{th}$ and $75^{th}$ percentiles, with the whiskers plotted between the hinges to the largest and smallest values at most 1.5x the interquartile range from their respective hinges. Accuracy metrics shown for each stimulation are mean 5-fold cross-validation accuracy score, and accuracy scores of null models generated with permuted group labels. For each metric, 100 trials of 5-fold cross validation were run. Statistical significance was determined via pairwise Mann-Whitney U tests, (**** $p < 0.0001$). PLS-DA scores plots for classifier models built on lectin binding data to distinguish unstimulated conditions to C) IL17 stimulation, D) IL4 stimulations, E) TNFa stimulation, F) IL6 stimulation, G) IL10 stimulation. Scores plot ellipses represent a 5 group confidence interval. H) PLS-DA scores plot for classifier model built on lectin binding data to distinguish samples from Bimm pool 1 and Bimm pool 2. Scores plot ellipses represent a 5 group confidence interval.

**Table 1.** High and low cytokine concentrations for 8 cytokine stimulation conditions.

| Cytokine | High concentration (ng/mL) | Low concentration (ng/mL) |
|---|---|---|
| IL-4 | 50 | 20 |
| IL-6 | 50 | 15 |
| IL-10 | 50 | 25 |
| IL-17 | 2.5 | 0.5 |
| TNFa | 50 | 5 |
| IFNg | 50 | 20 |
| APRIL | 12.5 | 3.13 |
| BAFF | 50 | 5 |

**Table 2.** Panel of 42 glycogenes, adapted from Nguyen et al. *Sci Rep.* (2021)

| Group | Gene Abbreviation | Gene |
|---|---|---|
| Nucleotide sugar synthesis | GALE | UDP-galactose 4-epimerase |
| Nucleotide sugar synthesis | GNE | UDP-N-acetylglucosamine-2 epimerase (Human) |
| Nucleotide sugar synthesis | NANS | Sialic acid synthase |
| Nucleotide sugar synthesis | NANP | N-acetylneuraminic acid phosphatase |
| Nucleotide sugar synthesis | CMAS | Cytidine monophospho-sialic acid synthase |
| Nucleotide sugar synthesis | CST | CMP-sialic acid transporter |
| Nucleotide sugar transporter | UGT | UDP-galactose transporter |
| Nucleotide sugar transporter | UGNT | UDP-N-acetylglucosamine transporter |
| Nucleotide sugar transporter | GFT | GDP-fucose transporter |
| Nucleotide sugar transporter | GANC | Neutral α-glucosidase C |
| Nucleotide sugar transporter | MANEA | Endo-α mannosidase |
| Nucleotide sugar transporter | MAN1A1 | Mannosyl-oligosaccharide 1,2-α-mannosidase IA |
| Nucleotide sugar transporter | MAN1B | Mannosyl-oligosaccharide 1,2-α-mannosidase IB |
| Glycan- processing glycosidase | MAN1C1 | Mannosyl-oligosaccharide1,2-α-mannosidase IC (isoform 1) |
| Glycan- processing glycosidase | MAN2A2 | α-mannosidase 2A member 2 |
| Glycan- processing glycosidase | MAN2B1 | α-mannosidase, class 2B, member 1 |
| Glycan- processing glycosidase | MAN2B2 | α-mannosidase, class 2B, member 2 |
| Glycan- processing glycosidase | MAN2C1 | α-mannosidase, class 2C, member 1 |
| Glycan- processing glycosidase | MGAT1 | α-1,3-mannosyl-glycoprotein 2-β-N-acetylglucosaminyltransferase |
| Glycan- processing glycosidase | MGAT2 | α-1,6-mannosyl-glycoprotein 2-β-N-acetylglucosaminyltransferase |
| Glycan- processing glycosidase | MGAT3 | β-1,4-mannosyl-glycoprotein 4-β-N-acetylglucosaminyltransferase |
| N-Glycan chain extension | MGAT4A | α-1,3-mannosyl-glycoprotein 4-β-N-acetylglucosaminyltransferase A |

| | | |
|---|---|---|
| N-Glycan chain extension | MGAT4B | α-1,3-mannosyl-glycoprotein 4-β-N-acetylglucosaminyltransferase B |
| N-Glycan chain extension | MGAT4C | α-1,3-mannosyl-glycoprotein 4-β-N-acetylglucosaminyltransferase C |
| N-Glycan chain extension | MGAT5 | α-1,6-mannosyl-glycoprotein 6-β-N-acetylglucosaminyltransferase |
| N-Glycan chain extension | MGAT5B | α-1,6-mannosyl-glycoprotein 6-β-N-acetylglucosaminyltransferase B |
| N-Glycan chain extension | B4GALT1 | β-1,4-Galactosyltransferase 1 |
| N-Glycan chain extension | B4GALT2 | β-1,4-Galactosyltransferase 2 |
| N-Glycan chain extension | B4GALT3 | β-1,4-Galactosyltransferase 3 |
| Galactosylation | B4GALT4 | β-1,4-Galactosyltransferase 4 |
| Galactosylation | B4GALT5 | β-1,4-Galactosyltransferase 5 |
| Galactosylation | B4GALT6 | β-1,4-Galactosyltransferase 6 |
| Galactosylation | B4GALT7 | β-1,4-Galactosyltransferase 7 |
| Galactosylation | ST3GAL1 | β-Galactoside -α-2,3-sialyltransferase 1 |
| Galactosylation | ST3GAL2 | β-Galactoside -α-2,3-sialyltransferase 2 |
| Galactosylation | ST3GAL3 | β-Galactoside -α-2,3-sialyltransferase 3 |
| Sialylation | ST3GAL4 | β-Galactoside -α-2,3-sialyltransferase 4 |
| Sialylation | ST3GAL5 | β-Galactoside -α-2,3-sialyltransferase 5 |
| Sialylation | ST3GAL6 | β-Galactoside -α-2,3-sialyltransferase 6 |
| Sialylation | ST6GAL1 | β-Galactoside -α-2,6-sialyltransferase 1 |
| Sialylation | ST6GAL2 | β-Galactoside -α-2,6-sialyltransferase 2 |
| Fucosylation | FUT8 | α-1,6-Fucosyltransferase |

**Table 3.** Lectins used in Enzyme-Linked Lectin Assay (ELLA)

| Lectin | Glycan target | Source, Catalog Number | Concentration for ELLA (μg/mL) |
|---|---|---|---|
| Sambucus Nigra Agglutinin (SNA) | α2-6-sialylated LacNAc | Vector Laboratories, B-1305-2 | 0.1 |
| Ricinus Communis Agglutinin I (RCA) | Terminal Type 2 LacNAc | Vector Laboratories, B-1085-1 | 1 |
| Lens Culinaris Agglutinin (LCA) | Core fucose | Vector Laboratories | 10 |
| Maackia Amurensis-I (MAL-I) | α2-3-sialylated LacNAc | Vector Laboratories, B-1315-2 | 10 |
| Phaseolus Vulgaris Erythroagglutinin (PHA-E) | Bisecting GlcNAc with Type 2 LacNAc | Vector Laboratories, B-1125-2 | 1 |
| Wheat Germ Agglutinin (WGA) | Terminal GlcNAcβ<br>Terminal GlcNAcα<br>Terminal NAc containing glycans | Vector Laboratories, B-1025-5 | 1 |
| Datura Stramonium Lectin (DSL) | Chitin<br>Type 2 poly LacNAc | Vector Laboratories, B-1185-2 | 1 |
| Phaseolus Vulgaris Leucoagglutinin (PHA-L) | β1-6 branched N-glycans | Vector Laboratories, B-1115-2 | 1 |
